# Engineered Reproductively Isolated Species Drive Reversible Population Replacement

**DOI:** 10.1101/2020.08.09.242982

**Authors:** Anna Buchman, Isaiah Shriner, Ting Yang, Junru Liu, Igor Antoshechkin, John M. Marshall, Michael W. Perry, Omar S. Akbari

## Abstract

Engineered reproductive species barriers are useful for impeding gene flow and driving desirable genes into wild populations in a reversible threshold-dependent manner. However, methods to generate synthetic barriers are lacking in advanced eukaryotes. To overcome this challenge, we engineered SPECIES (<u>S</u>ynthetic <u>P</u>ostzygotic barriers <u>E</u>xploiting <u>C</u>RISPR-based <u>I</u>ncompatibilities for <u>E</u>ngineering <u>S</u>pecies) to generate postzygotic reproductive barriers. Using this approach, we engineer multiple reproductively isolated SPECIES and demonstrate their threshold-dependent gene drive capabilities in *D. melanogaster*. Given the near-universal functionality of CRISPR tools, this approach should be portable to many species, including insect disease vectors in which confinable gene drives could be of great practical utility.

**One Sentence Summary:** Synthetically engineered SPECIES drive confinable population replacement.

## Results

Species that are reproductively incompatible with, but otherwise identical to, wild counterparts can be engineered via the insertion of reproductive barriers. Such barriers could be used for ecosystem engineering or pest and vector control ^1^. Previous attempts have engineered synthetic species barriers via genetic recoding in bacteria ^2^ and yeast ^3^, though this is likely infeasible in multicellular organisms. A reproductively isolated strain of *D. melanogaster* was generated with preexisting transgenes and recessive mutations ^4^, though elevated fitness costs prevented utility of this approach. CRISPR-based genome editing and transcriptional transactivation (CRISPRa) strategies exist ^1,5^, though fail to achieve complete reproductive isolation due to either escape mutants or incomplete lethality.

Here, we describe the development of synthetic reproductive barriers in *D. melanogaster* using an approach we term “SPECIES” (<u>S</u>ynthetic <u>P</u>ostzygotic barriers <u>E</u>xploiting <u>C</u>RISPR-based <u>I</u>ncompatibilities for <u>E</u>ngineering <u>S</u>pecies) with similarities to a technique also recently described^6^. To engineer SPECIES, we use a nuclease-deficient deactivated Cas9 (dCas9) protein fused to a transactivation domain, which recruits transcriptional machinery to the site of single guide RNA (sgRNA) binding. These sgRNAs target the promoter region of an endogenous gene essential for development/viability to generate lethality mediated by dCas9 mediated overexpression. This lethality is rescued in engineered, but not WT, individuals via mutation(s) of the sgRNA target site(s), preventing dCas9 binding, and subsequent lethal overexpression, without interfering with target gene function (Fig. 1A).

**Figure 1.**
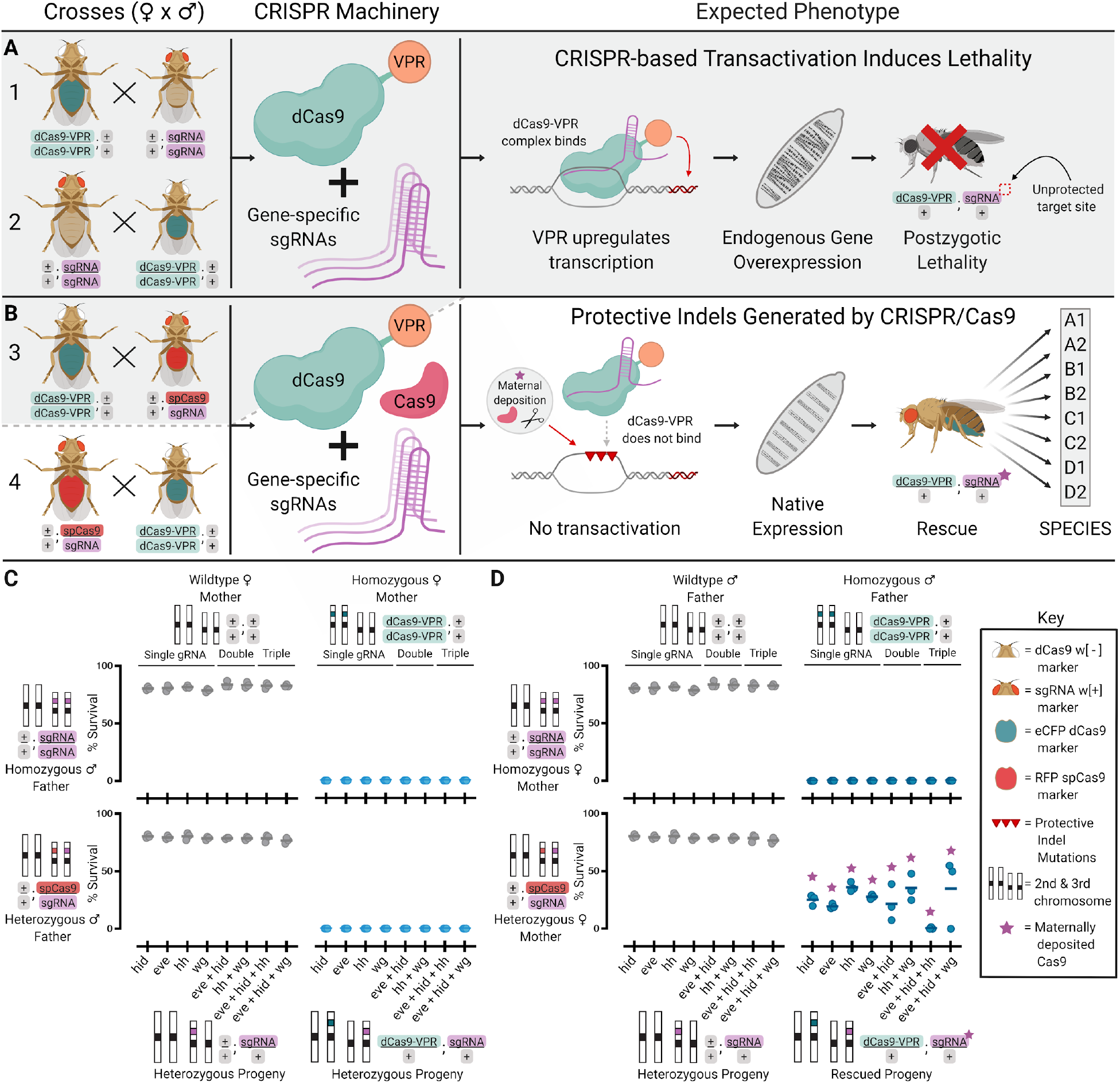
Development of Synthetic Reproductive Barriers. (**A**) Homozygous dCas9-VPR flies were crossed bidirectionally to homozygous sgRNA flies. Shaded grey background indicates expected lethal crosses. (**B**) Homozygous dCas9-VPR individuals were crossed bidirectionally to heterozygous spCas9/sgRNA individuals. Surviving individuals were repeatedly inbred to generate SPECIES. (**C,D**) Plots depicting % progeny survival from the crosses depicted in A and B (Tables S1, S2). All crosses were performed in triplicate. Horizontal bars indicate mean.

We engineered flies expressing a dCas9 activator domain fusion (dCas9-VPR) and evaluated whether these transgenes could drive lethal target overexpression using CRISPRa sgRNA lines each targeting the promoter region of one of four important developmental genes (*eve*, *hid*, *hh,* and *wg)* ^5,7,8^ (Fig. S1). Zygotic dCas9-VPR expression did not cause noticeable toxicity and achieved 100% lethality in individuals also expressing sgRNAs targeting one of the genes (Fig. 1B-D, Table S1). Interestingly, this lethality could only be rescued when homozygous dCas9-VPR-expressing fathers were crossed to heterozygous Cas9; sgRNA mothers (Fig. 1D, Table S2). With this cross, mothers provided both indel mutations in the promoter region of the target genes while simultaneously depositing sgRNA/Cas9 into all embryos, which mutated the inherited paternal copy of the target sites.

Importantly, the inherited sgRNA/dCas9-VPR transgenes forced a bottleneck that selected for protective indels that did not compromise the expression level of the target gene, providing embryonic rescue and survival (Fig. 1C,D, S2, Table S1). However, in crosses of WT and heterozygous dCas9-VPR; sgRNA individuals harboring the protective indel mutations, a very small fraction of progeny inheriting the sgRNA and dCas9-VPR transgenes survived, even with an inherited WT copy of the target site (Fig. S2A). Furthermore, homozygous dCas9-VPR/dCas9-VPR; sgRNA/sgRNA individuals outcrossed to WT produced viable progeny, indicating homozygous “rescued” flies were not completely reproductively isolated from their WT counterparts (Fig. S2), a feature also previously demonstrated by others ^5^.

To overcome the incomplete isolation from single-gene overexpression, we tested multiplexed overexpression by engineering flies that simultaneously expressed sgRNAs targeting two or more genes (*eve + hid; eve + hid + hh; eve + hid + wg;* and *hh + wg*; Fig. S1, S2). Crossing these to the dCas9-VPR flies resulted in complete progeny lethality (Fig. 1C, Table S2), suggesting that heterozygosity for a WT allele and an allele with a selected indel is lethal.

**Figure 2.**
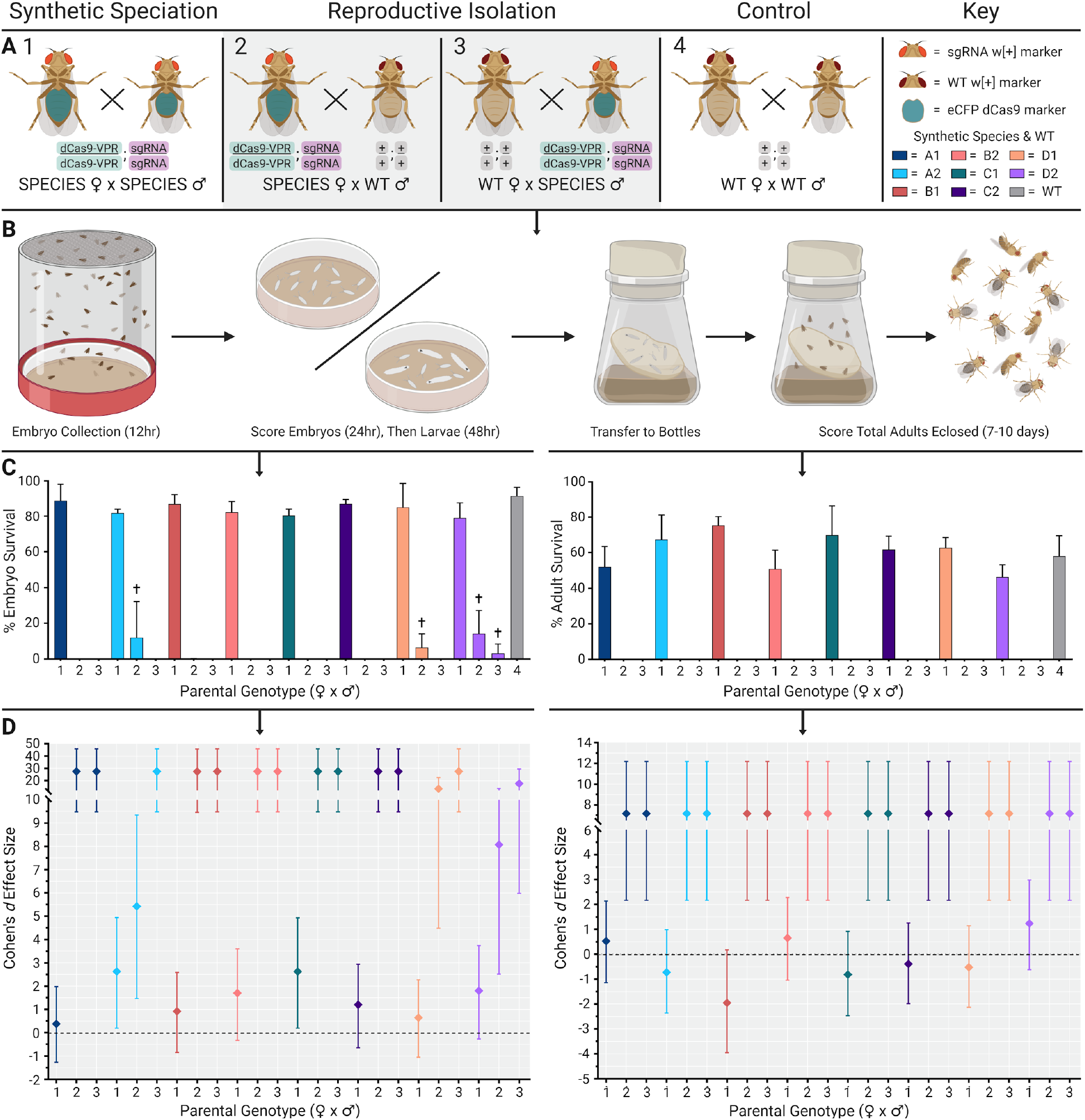
Embryo and adult viability determination. (**A**) Crosses used to determine embryo and adult viability. (**1**) Homozygous dCas9-VPR;sgRNA “SPECIES” females crossed to homozygous dCas9-VPR;sgRNA SPECIES males. (**2**) Homozygous dCas9-VPR;sgRNA SPECIES females crossed to wildtype (WT) males. (**3**) WT females crossed to homozygous dCas9-VPR;sgRNA SPECIES males. Grey shaded background indicates expected lethal crosses. (**4**) WT females crossed to WT males. (**B**) Schematic detailing the methods of determining embryo and adult survival compared to WT. (**C**) % embryo and adult survival was calculated and plotted. The number below each bar indicates cross type (1-4). ✝ indicates that embryos did not survive past L1/L2 stages. Unpaired t-tests were performed for each SPECIES compared to WT (Table S3). Error bars indicate SD. (**D**) Embryo and adult Cohen’s *d* effect sizes compared to WT. Error bars indicate 95% CI.

With the selective bottleneck genetic crossing scheme, crosses with multiplexed sgRNA/Cas9-expressing mothers rescued heterozygous dCas9; sgRNA animals through the introduction of indel mutations (Fig. S4). Some fitness costs can be seen, at least as inferred from fertility and survivorship, can be detected in many of the SPECIES strains (Table S3).

Moreover, in contrast to the single-target sgRNA lines, heterozygote progeny from dCas9-VPR; multiplexed sgRNA individuals crossed to WT were 100% lethal, suggesting that inheriting one WT copy of each target site from the WT parent ensured lethality (Fig. S2).

To generate reproductively isolated SPECIES, multiple generations (>5) of dCas9-VPR; sgRNA “rescued” individuals were intercrossed, resulting in homozygous stocks representing eight isolated SPECIES (A1-D2). Each SPECIES was reproductively incompatible with WT (Fig. S2, S3) and harbored the expected indels at the target sites (Fig. S4). Bidirectional outcrosses of all eight SPECIES to WT, or to a different SPECIES with varying target genes, demonstrated 100% reproductive isolation, indicating the creation of several independent barriers to sexual reproduction (Fig. 3, S5, Table S3, Video S1). Additional crosses between SPECIES and genetically diverse stocks from five different continents also demonstrated 100% reproductively isolation (Table S19). We next determined the extent of target gene overexpression when outcrossed to WT by visualizing overexpression in embryos via antibody stain, and we evaluated the effect of misexpression on development using cuticle preps of late embryos and young larvae. We observed target gene overexpression at embryonic stages and segment polarity defects in larvae when the SPECIES lines were mated to WT but not when self-crossed (Fig. 3A-C). To quantify the extent of target-gene overexpression and to measure possible global gene misexpression, we performed transcriptome-wide expression profiling (Table S5). We quantified RNA-expression profiles for all samples (Table S6), including genes that were expressed from our constructs (Table S7). From this analysis, we found significant target gene overexpression (up to 48-fold) in the progeny generated from SPECIES and WT crosses but not in the progeny from SPECIES intercrosses (Fig. 3D, S6, S7, Tables S6-S12, S16, S18).

**Figure 3.**
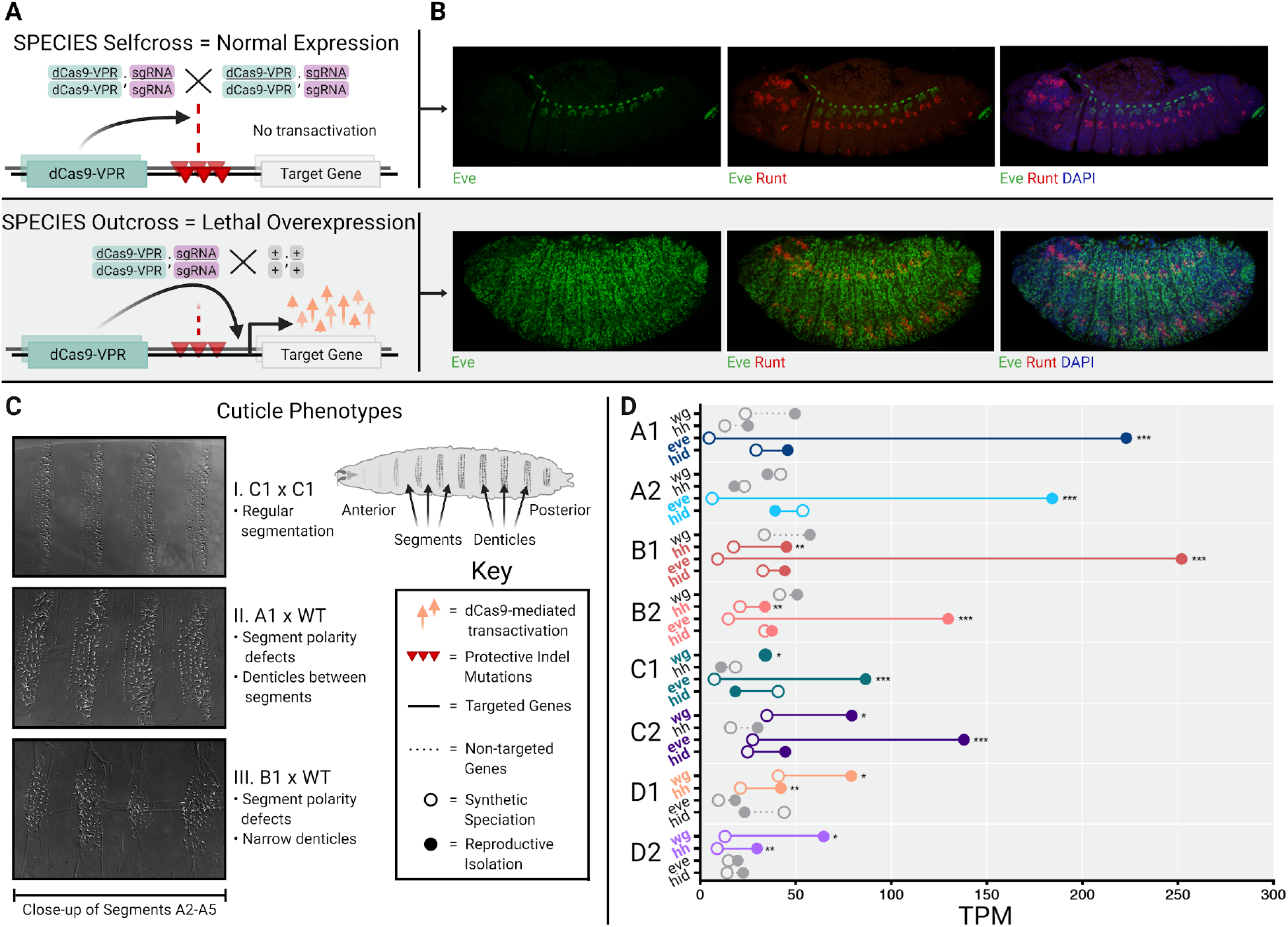
Visualization and Quantification of Target Gene Overexpression. (**A**) Schematic demonstrating reproductive barriers. The homozygous progeny of intercrossed SPECIES contain two copies of protected indel mutations, preventing lethal overexpression. The heterozygous progeny of SPECIES outcrossed to WT inherit only one copy of the protected indel mutations, which cannot prevent lethal overexpression of the target genes. Grey shaded background indicates expected lethality. (**B**) Antibody stains for Eve (green) and Runt (red) in stage-13 embryos. DAPI (blue) stains all nuclei. Top row shows A1xA1. Bottom row shows A1xWT SPECIES outcross and Eve overexpression. (**C**) Cuticle preparations showing denticle belts in self cross vs. outcross. (**D**) Normalized embryo RNAseq data (transcripts per kilobase million, TPM) for each species either self-crossed (open circles) or outcrossed to WT (closed circles), indicating expected target gene overexpression (colored) (Table S6, S7). Unpaired t-tests were performed for each gene, comparing self-cross of the SPECIES targeting the gene to the outcross data of those same SPECIES (*P < 0.05, **P < 0.01, ***P < 0.001, and ****P < 0.0001) (Table S16).

To assess whether the SPECIES were capable of reversible WT population replacement via gene drive, we conducted population studies using bottles at various release thresholds using one representative SPECIES, A1 (Fig. 4). Releases of A1 individuals at a population frequency of 70% resulted in this SPECIES replacing the WT population in two of six replicates (Tables S13, S14). Population replacement also occurred in three of four replicates of A1 releases at a frequency of 80% and in one of one replicate at a 90% release frequency. However, a release frequency of 50% resulted in elimination of the A1 strain in three of three replicates (Fig. 4, Tables S13, S14).

**Figure 4.**
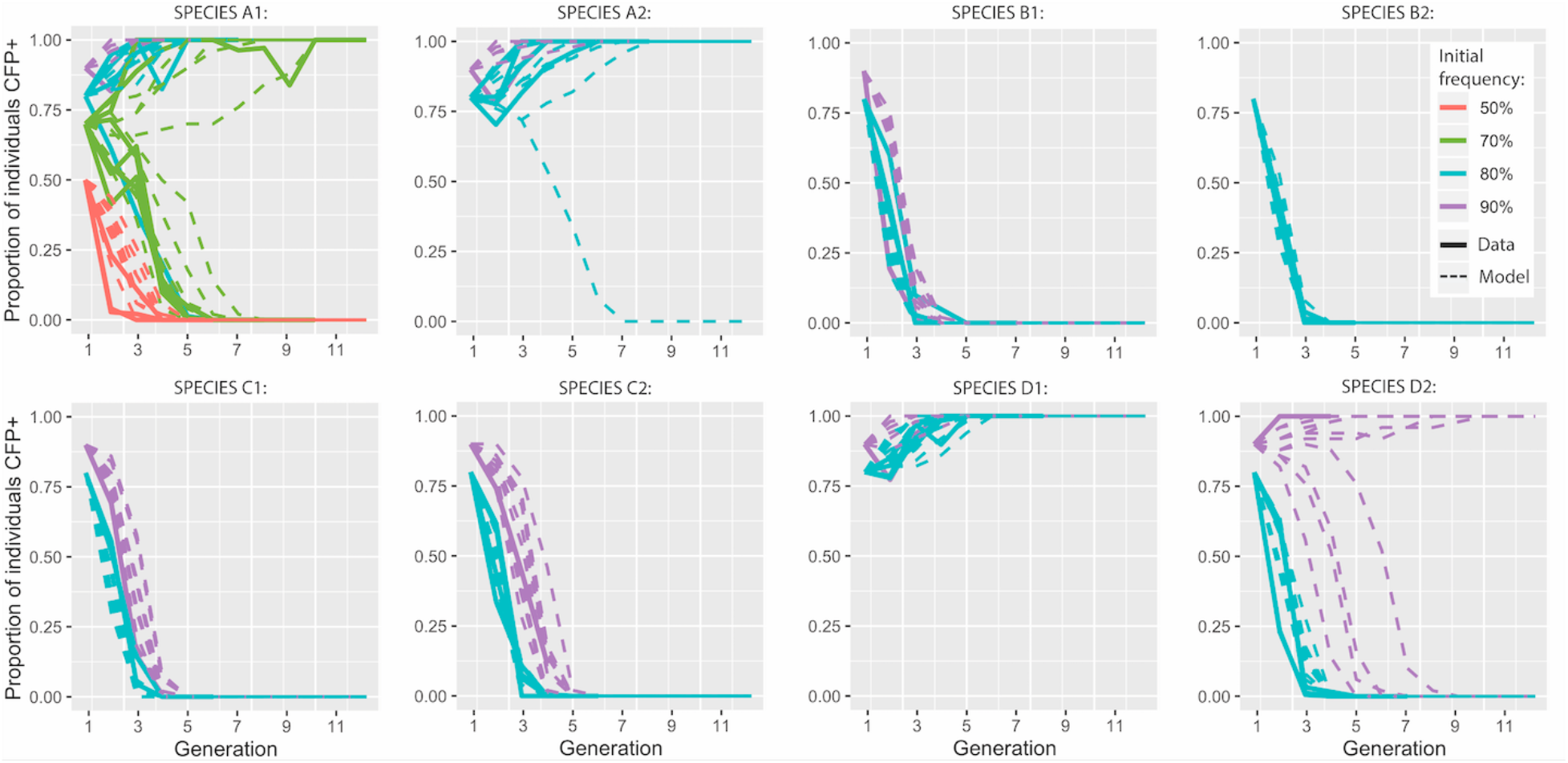
Population Experiments and Model Fits for Eight SPECIES Demonstrating Threshold-dependent Population Replacement. Population experiments mated SPECIES individuals with WT individuals, producing first generation SPECIES population frequencies of 50%, 70%, 80%, and 90% for system A1; 80% and 90% for systems A2, B1, C1, C2, D1, and D2; and 90% for system B2. Results are shown as solid lines, while fitted model predictions are dashed lines. Observed data are consistent with fitness costs of SPECIES strains relative to WT (Table S17). Ten stochastic model predictions are shown for each release frequency, assuming a population size of 50. Proportion of individuals CFP+ represents the percentage of SPECIES individuals at each generation.

To characterize the population dynamics observed in the population studies, we fitted a mathematical model to the observed data, incorporating a fitness cost for reproductively isolated individuals relative to WT individuals. The A1 strain was estimated to have a strong relative fitness cost of 34.84% (95% credible interval [CrI]: 34.82–34.87%), producing a threshold frequency of ~61%, which corresponds to what was observed in the population studies. Of the seven other SPECIES characterized, two consistently led to population replacement at a release frequency of 80% (A2 and D1), and three led to population replacement at a release frequency of 90% (A2, D1 and D2) (Fig. 4), with increased threshold frequencies corresponding to increased fitness costs for all SPECIES (Table S17). This suggests that population replacement via gene drive would theoretically occur when the release of SPECIES individuals exceeded a critical threshold frequency in the population ^9,10^, the value of which depends on the fitness of the synthesized strain relative to the WT strain.

## Discussion

Altogether, we demonstrate that our SPECIES approach can be exploited to build reproductive barriers that could drive genes through a population in a reversible manner. The SPECIES approach is advantageous over other developed technologies, as dCas9-mediated overexpression does not rely on any SPECIES-specific mode of incompatibility; instead, it is programmable to virtually any suitable target gene and thus should be amenable to most sexually-reproducing organisms. Additionally, the “stacked” genetic barriers using more than one sgRNA may reduce both failure and the chances of resistance via target site mutation. Once a basic engineering toolkit is constructed, it can build multiple SPECIES that are reproductively isolated. Furthermore, SPECIES underdominant systems are preferable to other underdominant systems, such as translocations, for local population replacement since they can tolerate higher fitness costs, spread more quickly, lead to less contamination of neighboring populations, and are more resilient to elimination due to immigration of wild-types (Figs. 5, 6). Moreover, while the SPECIES approach was recently demonstrated in flies ^6^, our study complements that work and took the approach a step further. For example, (i) we and others ^5^ demonstrate that single gene targeting is insufficient to generate SPECIES and therefore exploit multiplexing of the target genes (2-3 targets per SPECIES) which also increases the evolutionary stability; (ii) we exploit a genetic crossing scheme that forces a genetic bottleneck which streamlines the genetics and enables the organism to select for non-deleterious indels. This crossing scheme enabled us to multiplex our targets and also may also serve important for generating SPECIES in polyploids (eg. fish and plants); (iii) our study also demonstrated the majority-wins threshold dependent gene drive capabilities of SPECIES and provides detailed modeling to support this application.

**Figure 5.**
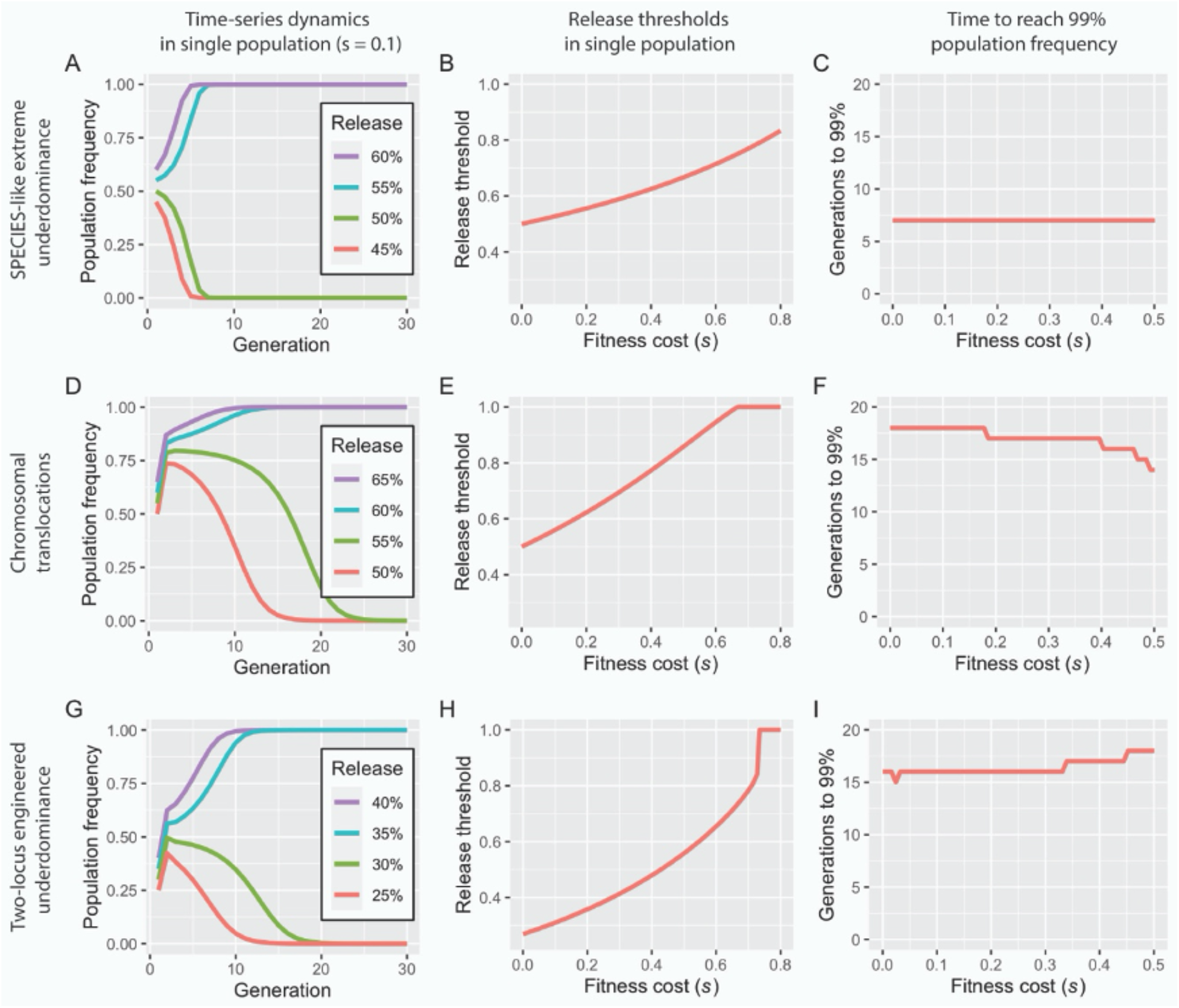
Population dynamics of underdominant systems in a single population. (**A**) Discrete generation model of a SPECIES-like extreme underdominant system with a fitness cost of 10% released at population frequencies of 45-60%. A release threshold is apparent between 50-55% (simulations confirm a threshold of 52.8%). (**B**) The release threshold increases with fitness cost, from 50% for no fitness cost, to 83.3% for an 80% fitness cost. (**C**) Extreme underdominant systems spread quickly, reaching a population frequency of 99% (including heterozygotes and homozygotes) within 7 generations of a release at a frequency 1% above the release threshold for fitness costs between 0-50%. (**D**) Discrete generation model of reciprocal chromosomal translocations with a fitness cost of 10% released at population frequencies of 50-65%. A release threshold is apparent between 55-60% (simulations confirm a threshold of 56.1%). (**E**) The release threshold increases with fitness cost, from 50% for no fitness cost, to 62.4% for a 20% fitness cost. Translocations cannot spread for fitness costs greater than 66.3%. (**F**) Translocations spread less quickly, reaching a population frequency of 99% (including heterozygotes and homozygotes) within 14-18 generations of a release at a frequency 1% above the release threshold for fitness costs between 0-50%. (**G**) Discrete generation model of two-locus engineered underdominance with a fitness cost of 10% released at population frequencies of 25-40%. A release threshold is apparent between 30-35% (simulations confirm a threshold of 31.2%). (**H**) The release threshold increases with fitness cost, from 26.9% for no fitness cost, to 35.9% for a 20% fitness cost. Two-locus engineered underdominance cannot spread for fitness costs greater than 72.7%. (**I**) Two-locus engineered underdominant systems also spread less quickly, reaching a population frequency of 99% (including heterozygotes and homozygotes) within 15-18 generations of a release at a frequency 1% above the release threshold and for fitness costs between 0-50%.

**Figure 6.**
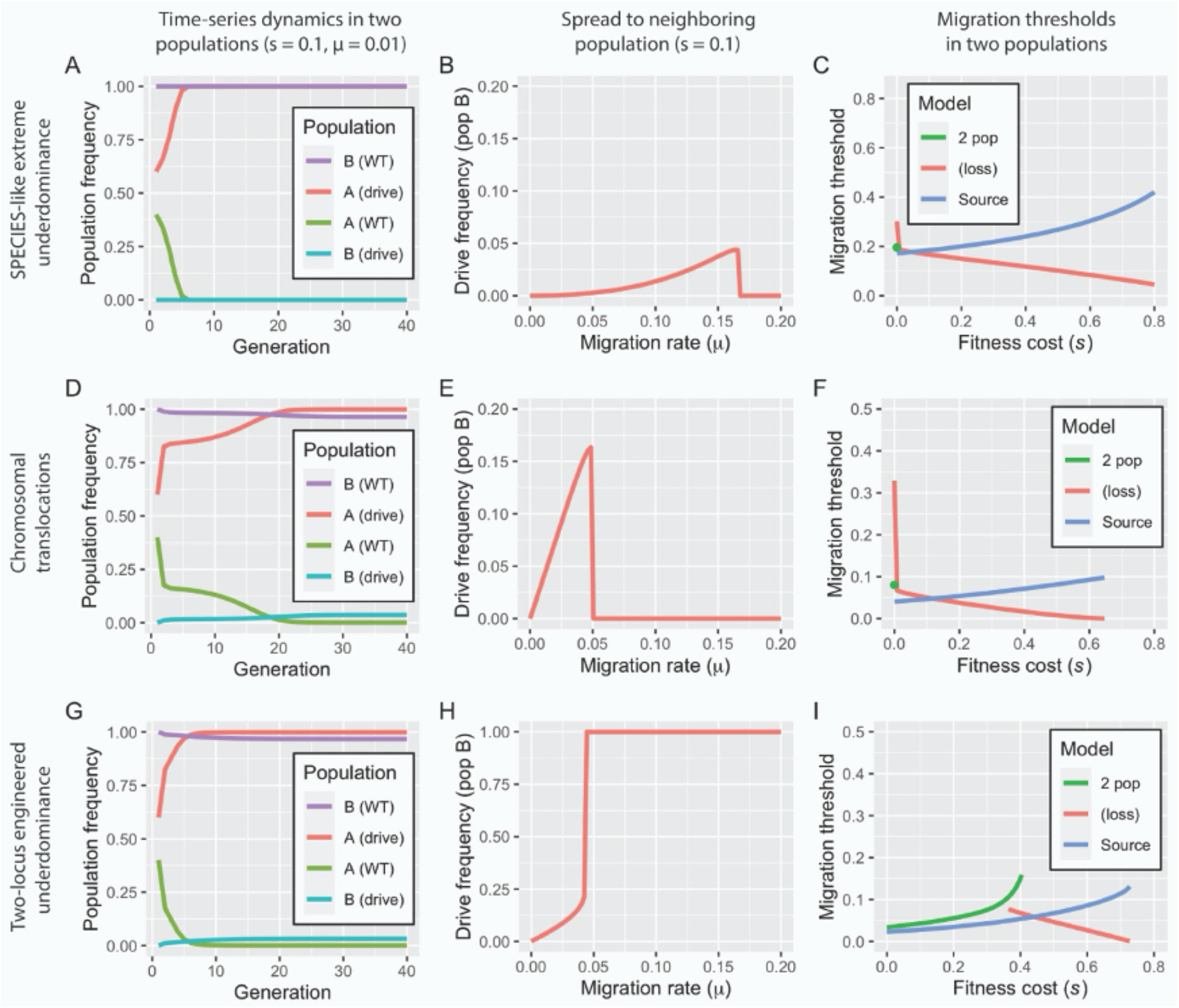
Population dynamics of underdominant systems in two populations. (**A**) Discrete generation model of a SPECIES-like extreme underdominant system released at 60% in population A and initially absent from population B. Population A exchanges migrants with population B at a rate of 1% per individual per generation. For a fitness cost of 10%, the system reaches near-fixation in population A within 7 generations but only spreads to 0.01% in population B. (**B**) As the migration rate increases, the SPECIES system reaches a higher frequency in population B, exceeding 4%; however, for migration rates above 16.6% per individual per generation, it is eliminated from both populations through dilution of population A with wild-types from population B. (**C**) For the two-population model, there is a migration threshold below which the construct fixes in population A and persists at a low level in population B and above which it is lost in both populations. For the source model, extreme underdominance displays threshold behavior with respect to migration rate. (**D**) Discrete generation model of reciprocal chromosomal translocations released at 60% in population A and initially absent from population B. Population A exchanges migrants with population B at a rate of 1% per individual per generation. For a fitness cost of 10%, the system reaches near-fixation in population A within 22 generations and spreads to 3.6% in population B. (**E**) As the migration rate increases, the translocations reach a higher frequency in population B, exceeding 15%; however, for migration rates above 5.0% per individual per generation, they are eliminated from both populations through dilution of population A with wild-types from population B. (**F**) For the two-population model, there is a migration threshold below which translocations fix in population A and persist at a low level in population B, and above which they are lost in both populations. For the source model, translocations display threshold behavior with respect to migration rate. (**G**) Discrete generation model of two-locus engineered underdominance released at 60% in population A and initially absent from population B. Population A exchanges migrants with population B at a rate of 1% per individual per generation. For a fitness cost of 10%, the system reaches near-fixation in population A within 8 generations and spreads to 3.2% in population B. (H) As the migration rate increases, the system reaches a higher frequency in population B, exceeding 21.2%; however, for migration rates above 4.2% per individual per generation, the system becomes fixed in both populations. (I) Two-locus engineered underdominance displays threshold behavior with respect to migration rate.

Bringing the SPECIES approach to other organisms will require integrating a cassette containing dCas9-VPR and sgRNAs into the genome of interest, which is quickly becoming more tractable with CRISPR-based tools. This approach will also require genome characterization to provide candidate sgRNA sites in promoter sequences near genes of interest as well as species-specific testing for overexpression levels and dCas9-VPR toxicity, which may contribute to observed fitness costs ^11^. Regardless, it should be possible to implement SPECIES in organisms of medical or agricultural interest, such as mosquitoes. This becomes even more interesting considering the gene drive function of SPECIES provides greater control and confinement via threshold dependence ^12–14^ as well as a reversibility via WT release ^15^ and protection from the evolution of resistance via multiplexing, features not found in all gene drives ^13,16^. While release requirements for threshold-dependent systems are large due to fitness costs, they are an order of magnitude less than those routinely carried out for insect suppression programs ^17^ and spatially-explicit models suggest that local population replacement could be achieved with 1:1 releases over several weeks ^18^. Overall, SPECIES demonstrates a platform for the possible safe control or modification of pest and disease vector populations that impose significant burdens on humanity.

## Materials and Methods

### Plasmid design and assembly

To assemble plasmid OA-986A, the base vector used for generating dCas9-expressing plasmids, several components were cloned into the piggyBac plasmid pBac[3xP3-DsRed] ^19^ using Gibson assembly/EA cloning ^20^. pBac[3xP3-DsRed] was digested with BstBI and NotI, and the following components were cloned in with EA cloning: an *attP* sequence amplified from plasmid M{3xP3-RFP attP} ^21^ with primers 986.C1 and 986.C2, a p10 3’UTR fragment amplified from Addgene plasmid #100580 ^19^ with primers 986.C3 and 986.C4, an opie2 promoter fragment amplified from translocation plasmid B ^15^ using primers 986.C5 and 986.C6, and an eCFP marker amplified from Addgene plasmid #47917 using primers 986.C7 and 986.C8. The resulting plasmid was then digested with PacI, and the following components were cloned in to generate the final dCas9-expressing vectors: the *Ubiquitin-63E* promoter fragment amplified with primers 986.C9 and 986.C10 from *D. melanogaster* genomic DNA and a dCas9-VPR fragment amplified from Addgene plasmid #78898 ^22^ with primers 986.C11 and 986.C12 to generate plasmid OA-986B (Addgene #124999); the *bottleneck* promoter fragment amplified with primers 986.C13 and 986.C14 from *D. melanogaster* genomic DNA and a dCas9-VPR fragment amplified from Addgene plasmid #78898 ^22^ with primers 986.C15 and 986.C12 to generate plasmid OA-986C (Addgene #125000); the *Ubiquitin-63E* promoter fragment amplified with primers 986.C9 and 986.C16 from *D. melanogaster* genomic DNA and a dCas9-VP64 fragment amplified from Addgene plasmid #78897 ^22^ with primers 986.C17 and 986.C18 to generate plasmid OA-986D (Addgene #125001); and the *bottleneck* promoter fragment amplified with primers 986.C13 and 986.C19 from *D. melanogaster* genomic DNA and a dCas9-VP64 fragment amplified from Addgene plasmid #78897 ^22^ with primers 986.C20 and 986.C18 to generate plasmid OA-986E (Addgene #125002).

To assemble plasmids OA-1045A-E, the multiple-sgRNA containing vectors, several components were cloned into the multiple cloning site (MCS) of a plasmid ^21^ containing the *white* gene as a marker and an *attB*-docking site using Gibson assembly/EA cloning ^20^. First, the plasmid was digested with AsiSI and KpnI, and the following components were cloned in with EA cloning to generate base plasmid OA-1045: a *D. melanogaster* U6:3 promoter fragment sequence amplified from Addgene plasmid #49411 ^23^ with primers 1045.C1 and 1045.C2, and an sgRNA scaffold fragment amplified from Addgene plasmid #49411 ^23^ with primers 1045.C3 and 1045.C4. The resulting base plasmid was then used to clone final sgRNA plasmids OA-1045A through OA-1045E. To generate plasmid OA-1045A (Addgene #125003), plasmid OA-1045 was digested with AvrII; then, a fragment containing an 18-base-pair (bp) *even skipped (eve)* sgRNA target site ^5^, an sgRNA scaffold, a *D. melanogaster* U6:1 promoter fragment, and an 18-bp *head involution defective (hid)* sgRNA target site ^5^ was amplified from a custom gBlocks® Gene Fragment (Integrated DNA Technologies, Coralville, Iowa) with primers 1045.C5 and 1045.C6 and was cloned into the digested backbone using EA cloning. To generate plasmid OA-1045B (Addgene #125004), plasmid OA-1045A was digested with XbaI, and a fragment containing a *Gypsy* insulator, a *D. melanogaster* U6:1 promoter fragment driving expression of a first *hedgehog* (*hh)*-targeting sgRNA, and a *D. melanogaster* U6:3 promoter fragment driving expression of a second *hh*-targeting sgRNA amplified from plasmid pCFD4-*hh* ^8^ with primers 1045.C7 and 1045.C8 was cloned in using EA cloning. To generate plasmid OA-1045C (Addgene #125005), plasmid OA-1045A was digested with XbaI, and a fragment containing a *Gypsy* insulator, a *D. melanogaster* U6:1 promoter fragment driving expression of a first *wingless (wg)*-targeting sgRNA, and a *D. melanogaster* U6:3 promoter fragment driving expression of a second *wg*-targeting sgRNA amplified from plasmid pCFD4-*wg* ^8^ with primers 1045.C7 and 1045.C8 was cloned in using EA cloning. To generate plasmid OA-1045D (Addgene #125006), plasmid OA-1045 was digested with AscI and XbaI, and two fragments were cloned in using EA cloning: a first fragment containing a *D. melanogaster* U6:1 promoter fragment driving expression of a first *wg*-targeting sgRNA and a *D. melanogaster* U6:3 promoter fragment driving expression of a second *wg*-targeting sgRNA amplified from plasmid pCFD4-*wg* ^8^ with primers 1045.C9 and 1045.C10, and a second fragment containing a *Gypsy* insulator, a *D. melanogaster* U6:1 promoter fragment driving expression of a first *hh*-targeting sgRNA, and a *D. melanogaster* U6:3 promoter fragment driving expression of a second *hh*-targeting sgRNA amplified from plasmid pCFD4-*hh* ^8^ with primers 1045.C11 and 1045.C12. Finally, to generate plasmid OA-1045E (Addgene #125007), plasmid OA-1045 was digested with AvrII and NotI, and a fragment comprising *D. melanogaster* Gly tRNA-flanked sgRNAs^24^ targeting, from 5’ to 3’, *eve*, *hid*, and *hh* followed by a *D. melanogaster* U6:3 UTR that was amplified with primers 1045.C13 and 1045.C14 from a gene synthesized vector (GenScript, Piscataway, NJ) was cloned using EA cloning. All primers used for cloning are listed in Table S15.

### Fly culture and strains

Fly husbandry and crosses were performed under standard conditions at 25°C. Rainbow Transgenics (Camarillo, CA) carried out all of the fly injections. The fly strains used to generate dCas9-expressing lines were *attP* lines attP40w (Rainbow Transgenic Flies line; yw P{nos-phiC31\int.NLS}X;P{CaryP}attP40) and 8621 (BSC #8621; y[1] w[67c23]; P{y[+t7.7]=CaryP}attP1). The fly strains used to generate sgRNA-expressing lines were 86Fa (BSC #24486: y[1] M{vas-int.Dm}ZH-2A w[*]; M{3xP3-RFP.attP'}ZH-86Fa), 9732 (BSC #9732: y[1] w[1118]; PBac{y[+]-attP-9A}VK00013), and 8622 (BSC #8622: y[1]w[67c23]; P{y[+t7.7]=CaryP}attP2). For balancing chromosomes, fly stock BSC#39631 (w[*]; wg[Sp-1]/CyO; P{ry[+t7.2]=neoFRT}82B lsn[SS6]/TM6C, Sb[1]) was used. All lines were homozygous-viable.

### Generation and genetics of speciated stocks

Single sgRNA lines targeting *eve*, *hid, hh,* and *wg* were previously described ^5,8^. dCas9–VPR- and dCas9–VP64-expressing lines were generated via microinjection as described above; we were unable to generate a transgenic line that expressed dCas9-VPR ubiquitously, despite numerous attempts, suggesting that such expression was toxic. To test for the ability of all sgRNA lines to induce lethal overexpression (“killing”), 5 sgRNA males and 5 virgin females were separately crossed to 5 dCas9 line individuals of the opposite sex in single vials and were allowed to mate for 7 days. After 7 days, the parents were removed, and the vials were monitored for 7 additional days to assess whether viable larvae were present. No killing was observed in crosses of dCas9-VP64 expressing lines to any of the sgRNA-expressing lines (Table S1), consistent with previous observations ^8^. Complete killing was presumed when no larvae were present after 14 days. All experiments were done in biological triplicate.

To generate protective indel mutations, a previously described ^25^ *Ubiquitin*-Cas9 line (BSC #79005) was used. Briefly, 10 *Ubiquitin*-Cas9 virgin females were crossed to 10 sgRNA males, and virgin female and male progeny with both transgenes were selected and crossed to each other for at least three generations. Cas9/sgRNA trans-heterozygous virgins were then outcrossed in groups of 3–5 to homozygous attP40w *bnk*-dCas9-VPR males, and progeny containing both a sgRNA (identified by the presence of the w+ marker) and *bnk*-dCas9-VPR (identified by the presence of the opie2-eCFP marker) were isolated as “rescue” individuals that presumably carried protective indel mutations in the target promoter regions that prevented dCas9-induced overexpression (Table S1). To confirm the generation of indels, these flies were Sanger sequenced and crossed to each other, again in groups of 3–5, to establish “rescue” stocks.

These “rescue” crosses were also set up in the reverse direction, utilizing 3–5 homozygous attP40w *bnk*-dCas9-VPR females crossed to Cas9/sgRNA trans-heterozygous males, to determine whether maternal deposition of Cas9/sgRNAs is required for generating sufficient protective indel mutations to provide rescue of lethality. In particular, it was assumed that, if both copies of the targeted promoter needed to contain protective indel mutations to provide rescue, lack of maternally deposited Cas9/sgRNA (due to Cas9/sgRNA fathers being used) would lead to lack of “rescue” individuals, as all individuals inheriting the sgRNA and *bnk*-dCas9-VPR transgenes would still have one wildtype copy of the target promoter inherited from the mother available for targeting and would perish.

To further validate whether both copies of the targeted promoter needed to contain protective indel mutations to provide rescue from lethality, “rescue” individuals were also bidirectionally outcrossed in groups of 3–5 and in triplicate to homozygous attP40w *bnk*-dCas9-VPR individuals, and the resulting progeny were scored for the “rescue” phenotype. Here, it was presumed that the lack of transheterozygous sgRNA/*bnk*-dCas9-VPR progeny indicated that both copies of the targeted promoter needed to contain protective indel mutations to provide rescue and that the lack of such mutations in the promoter allele inherited from the homozygous attP40w *bnk*-dCas9-VPR parent led to lethality in the transheterozygous sgRNA/*bnk*-dCas9-VPR progeny. Here, too, such lethality was observed for crosses with multiple sgRNA transgenes but not for crosses with single sgRNA transgenes, suggesting that, in the case of the latter, one wildtype copy of the targeted promoter was not sufficient to lead to sgRNA/*bnk*-dCas9-VPR-induced lethality.

Double-homozygous speciated stocks were generated for all sgRNA combinations by crossing dCas9/sgRNA heterozygotes and identifying homozygous progeny by eye color (orange to dark red eyes for homozygotes versus yellow to light red eyes for heterozygotes, depending on sgRNA insertion site) and opie2-eCFP intensity. Putative double homozygous individuals were then outcrossed to w[1118] individuals of the opposite sex in groups of three per vial to test for reproductive isolation. Flies were allowed to mate and lay eggs for 7 days, and vials were checked daily for hatched embryos. Flies that failed to fruitfully mate with w[1118] were presumed to be reproductively isolated double homozygotes and were then crossed to putative double homozygotes of the opposite sex to generate a double homozygous, reproductively isolated stock for each sgRNA line.

### Assessment of reproductive isolation from various strains

To determine whether double-homozygous SPECIES lines were reproductively isolated from stocks that were genetically diverse, SPECIES individuals were outcrossed to various Global Diversity Lines (GDL) isolated from five different continents ^26^ and were used in previous work examining gene-drive function in different genetic contexts ^27,28^. Briefly, 5 double homozygous individuals from each SPECIES stock were outcrossed to 5 individuals of the opposite sex from each of five Global Diversity Lines (from Beijing, China; Ithaca, NY; the Netherlands; Tasmania, Australia; and Zimbabwe, Africa). All crosses were done bidirectionaly with respect to sex and in triplicate. Flies were allowed to mate and lay eggs for 7 days, and vials were checked daily for hatched embryos for the following 7 days. Lack of embryo hatching was presumed to indicate reproductive isolation (Table S19).

To assess reproductive isolation between double-homozygous SPECIES lines, inter-SPECIES crosses were performed by crossing 2 SPECIES virgin females with 2 SPECIES males from each strain. Flies were allowed to mate for 12–16 hours under standard conditions; following this period, the adult flies were removed and the embryos were counted (Table S4). The vials were aged at 26°C for 24 hours and subsequently scored for the number of hatched embryos (if reproductive isolation did not occur). The vials were then kept at 26°C for 7–10 days to ensure no pupae/adults emerged in instances of reproductive isolation or to count emerged adults in instances of incomplete reproductive isolation (Table S4).

### Embryo and adult viability determination

For embryo viability counts (Table S3), 3–4 day old adult virgin females were mated with males of the relevant genotypes for 2–3 days in glass vials supplemented with *Drosophila* medium and yeast paste. Following this period, the adults were transferred to an egg collection chamber containing a grape juice agar plate. Females were allowed to lay at 26°C for 12 hours, after which the adults were removed and the total number of embryos were scored. These embryos were kept on the agar surface at 26°C for 24 hours. The % survival was then determined by counting the number of unhatched embryos. One group of 100–300 embryos per cross was scored in each experiment, and each experiment was carried out in biological triplicate (total number of offspring scored is presented in Table S3). The results presented are averages from these three experiments. Embryo survival was normalized with respect to the % survival observed in parallel experiments carried out with the Oregon R wildtype strain, which was 91.66%. For adult fly counts (Table S3), the agar plates were transferred to 250 ml plastic bottles with *Drosophila* medium and kept at 26°C for 7–10 days. Following this period, the number of adults that emerged was scored. The percentages of adult survival presented are averages from each cross normalized with respect to the % survival observed in Oregon R, which was 58.04% (total number of offspring scored is presented in Table S3); all crosses were set in triplicate. Unpaired t-test statistical analyses using GraphPad Prism version 8.2.1 for Windows (GraphPad Software, San Diego CA, www.graphpad.com) were carried out for both embryo and adult fly counts to compare expected and observed values. For species crossed to themselves, significant differences were found in embryo survival for species A2 and C1 compared to WT crossed to itself (p values = 0.0322 and 0.0325, respectively) but not for any of the remaining six species. For species outcrossed to WT, significant differences were found for all bidirectional crosses of each species to WT when compared to WT crossed to itself. This significance was seen in experiments for both embryo and adult fly survival (*p* < 0.05). Cohen’s *d* effect sizes were calculated using Stata (StataCorp. 2019. *Stata Statistical Software: Release 16*. College Station, TX: StataCorp LLC). Figure icons were created with Biorender.com

### Population experiments

All genetic experiments were conducted in a high-security Arthropod Containment Level 2 (ACL2) barrier facility in accordance with protocols approved by the Institutional Biosafety Committee from University of California San Diego. Population experiments were carried out at 26°C, 12 hour/12 hour day/night cycle, with ambient humidity in 250 ml bottles containing Lewis medium supplemented with live, dry yeast. Starting populations for drive experiments included equal numbers of virgins and males of similar ages for each genotype. Speciated double homozygotes (dCas9/dCas9; +/+) were introduced at a population frequency of 80% for above-threshold drive experiments, and 50% for below-threshold drive experiments. Oregon R virgin females and males (+/+; +/+) of a similar age as the reproductively isolated individuals made up the remainder of the population. The total number of flies for each starting population was 100. All 50%, 70%, 80% population experiments were conducted in at least triplicate, with exception of SPECIES C1 seeded at 80% in which one of the replicates did not produce any progeny due to bacteria/mold contamination. Moreover, only one replicate was conducted for releases seeded at 90% for all species except B2 as again this replicate did not produce any progeny due to bacteria/mold contamination. In total we conducted 44 population cage experiments which were tracked for up to 11 generations. After being placed together, adult flies were removed after seven days. After another seven days, progeny were collected and the fraction of speciated double homozygous individuals was determined (Table S5). The progeny were then placed into a new bottle to initiate the next generation. No significant evidence of crowding in the 250 ml bottles was observed.

### Mathematical Modeling

We modeled SPECIES population dynamics under laboratory conditions assuming random mating and discrete generations. We considered a SPECIES allele, “T”, and a corresponding wildtype allele, “t”. Since heterozygotes for the SPECIES system are unviable, there are only two viable genotypes – TT and tt. We denote the proportion of organisms having the genotype TT at generation *k* by *p*_*k*_, and the proportion having the wildtype genotype at generation *k* by (1 − *p*_*k*_). By considering all possible mating pairs, and assuming a fitness cost for TT individuals relative to wildtype individuals, *s*, the frequency of TT individuals in the next generation is given by:

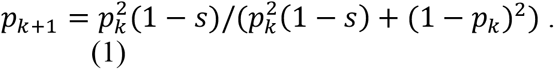

The threshold frequency is an unstable equilibrium that satisfies the condition:

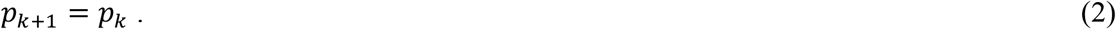

Substituting Equation 2 into Equation 1 and solving for *p*_*k*_, we find two stable equilibria (*p*_*k*_ = 0 and *p*_*k*_ = 1) and one unstable equilibrium (*p*_*k*_ = 1/(2 − *s*)). The latter represents the critical threshold frequency, above which the SPECIES system is more likely to spread to fixation than not, and below which it is more likely to be eliminated than not.

The likelihood of the population data for each SPECIES system was calculated by assuming a binomial distribution of wildtype (CFP-) and SPECIES (CFP+) individuals, and by using the model in Equation 1 to generate expected proportions for each fitness parameter value, *s*, i.e. by calculating the log-likelihood,

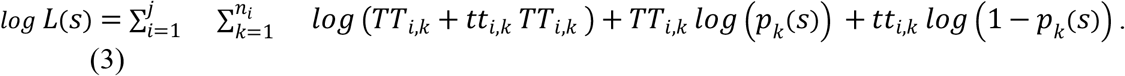

Here: i) *TT*_*i,k*_ and *tt*_*i,k*_ are the number of SPECIES (CFP+) and wildtype (CFP−) individuals at generation *k* in experiment *i*, respectively, ii) there are a total of *j* experiments for this SPECIES system, iii) the *i*th experiment is run for *ni* generations, and iv) the expected genotype frequencies are dependent on the fitness parameter, *s*. The initial condition for each experiment is specified by the data. Fitness parameters, including 95% credible intervals, were estimated using a Markov chain Monte Carlo (MCMC) sampling procedure.

The stochastic simulations in Fig. 4 were implemented by calculating expected genotype frequencies in the next generation according to Equation 1, and taking a binomial sample from a total of 50 individuals.

Comparative modeling of other underdominant systems is described in Marshall and Hay^29^. That paper uses the mathematical modeling framework described here in addition to two approaches for modeling migration: a) a “two-population model”, in which reciprocal movement occurs between the two connected populations; and b) a “source model”, in which the system is initially fixed in the source population, absent from the sink population, and one-way migration occurs from the source to sink population. In Marshall and Hay ^29^, population replacement and confinement dynamics are shown for: i) extreme underdominance (the SPECIES system modeled here), ii) reciprocal chromosomal translocations ^15^, iii) single-locus and two-locus engineered underdominance ^30^, iv) *Semele* ^31^, vi) inverse *Medea* ^31^, and vii) *Merea* (*Medea* with a recessive antidote). A range of parameter values are compared for each gene drive system, including fitness cost (*s*, varied between 0 and 30%) and migration rate (*m*, varied between 0 and 10% per individual per generation for both the source and two-population models) ^15^.

Results from that analysis suggest that SPECIES-like extreme underdominant systems fare well against other underdominance-based gene drive systems in terms of both confinement and persistence. The most direct comparison can be made to translocations ^15^, which also have a 50% release threshold in a single population and in the absence of a fitness cost. Considering a 5% fitness cost for both systems, they still have very similar release thresholds (51.3% for SPECIES-based underdominance c.f. 52.8% for translocations); however, for a two-population model with a migration rate of 1% per individual per generation, the SPECIES-based underdominant system spreads to only ~0.01% in the neighboring population, while the translocations spread to a much higher ~4.2% in the neighboring population ^29^. The migration rate at which the introduced system is lost due to inward migration of wild-types is also much higher for the SPECIES-based underdominant system (17.6% per individual per generation c.f. 5.8% for translocations, s= 0.05) ^29^. This suggests that SPECIES-like extreme underdominant systems are preferable to translocations for local population replacement since they lead to less contamination of neighboring populations and are less vulnerable to elimination due to inward migration.

Finally, were SPECIES-based underdominant systems to be implemented for local population replacement, strains would likely be used that would have much smaller fitness costs than those observed here (~30%). Despite that, results from Marshall and Hay ^15^ suggest the population dynamics of the SPECIES system are resilient in the face of these fitness costs. A SPECIES system with a fitness cost of 30% has a release threshold of 58.8%, which could be exceeded through weekly releases over several weeks. Furthermore, in a two-population model, the migration rate at which the SPECIES system would be lost due to inward migration of wild-types is 13.3% per individual per generation ^15^, which is greater than the movement rate observed between populations of *Anopheles gambiae* ^32^, the main mosquito vector of malaria in sub-Saharan Africa, and *Aedes aegypti* ^33^, the main mosquito vector of dengue, Zika and Chikungunya viruses.

### RNA sequencing for transcriptional activation analysis

Embryos were collected from the multiple speciated lines to assess transactivation in the embryo. Male speciated flies were crossed to Oregon R virgin females in glass vials supplemented with *Drosophila* medium and yeast paste and were incubated at 26°C for 72 hours. Following this period, the adult flies were transferred to collection chambers containing grape juice agar plates. The flies were allowed to lay for 4–5 hours, after which the embryos were aged for one hour and collected using a paintbrush. Afterwards, 30–50 embryos that were 5–6-hr old were collected, washed with ddH_2_O, and transferred to individual eppendorf tubes. The samples were flash frozen with liquid nitrogen and stored at −80°C. Intra-crosses for Oregon R, Cas-9, dCas-9, sgRNA, and speciated lines were also performed and collected as controls. Each sample was homogenized and processed using the Quick-Start Protocol of the miRNeasy Mini Kit (Qiagen, Hilden, DEU), followed by DNase treatment using the DNA-*free*™ Kit and protocol (Thermo Fisher Scientific, Waltham, MA, USA).

### RNA-seq library construction and sequencing

RNA integrity was assessed using the RNA 6000 Pico Kit for Bioanalyzer (Agilent Technologies #5067-1513), and mRNA was isolated from ~1 μg of total RNA using NEBNext Poly(A) mRNA Magnetic Isolation Module (NEB #E7490). RNA-seq libraries were constructed using the NEBNext Ultra II RNA Library Prep Kit for Illumina (NEB #E7770) following the manufacturer’s instructions. Briefly, mRNA was fragmented to an average size of 200 nt by incubating at 94°C for 15 min in the first strand buffer. cDNA was then synthesized using random primers and ProtoScript II Reverse Transcriptase followed by second strand synthesis using NEB Second Strand Synthesis Enzyme Mix. Resulting DNA fragments were end-repaired, dA tailed, and ligated to NEBNext hairpin adaptors (NEB #E7335). After ligation, adaptors were converted to the ‘Y’ shape by treating with USER enzyme, and DNA fragments were size selected using Agencourt AMPure XP beads (Beckman Coulter #A63880) to generate fragment sizes between 250 and 350 bp. Adaptor-ligated DNA was PCR amplified followed by AMPure XP bead clean up. Libraries were quantified using a Qubit dsDNA HS Kit (ThermoFisher Scientific #Q32854), and the size distribution was confirmed using a High Sensitivity DNA Kit for Bioanalyzer (Agilent Technologies #5067-4626). Libraries were sequenced on an Illumina HiSeq2500 in single read mode with the read length of 50 nt and sequencing depth of 20 million reads per library following the manufacturer's instructions. Base calls were performed with RTA 1.18.64 followed by conversion to FASTQ with bcl2fastq 1.8.4.

### Quantification and differential expression analysis

Reads were mapped to the *Drosophila melanogaster* genome (BDGP release 6, GenBank accession GCA_000001215.4) using STAR aligner ^34^ with the default parameters with the addition of the ‘--outFilterType BySJout’ filtering option and the ‘--sjdbGTFfile Drosophila_melanogaster.BDGP6.22.97.gtf’ GTF file downloaded from ENSEMBL. Expression levels were determined with featureCounts ^35^ using ‘-t exon -g gene_id -M -O --fraction’ options. Differential expression analyses between homozygous speciation stocks and corresponding heterozygotes outcrossed to wildtype females were performed with DESeq2 ^36^ using a two-factor design formula ‘design= ~ line + genotype’. Two independent lines per each target set (genotype) were used. MA plots [log2(FoldChange) vs log10(baseMean)] were generated with ggplot2 ^37^. All sequencing data can be accessed at NCBI SRA (study accession ID PRJNA578541, reviewer link: https://dataview.ncbi.nlm.nih.gov/object/PRJNA578541?reviewer=mnn67aeait5u7v231c1b8nh2vh).

### Immunohistochemistry

For antibody staining, embryos were collected overnight and then fixed and dechorionated using standard protocols ^38^. We used guinea pig anti-Runt polyclonal antibody (kindly provided by David Kosman and John Reinitz) at a concentration of 1:200 and mouse anti-Eve monoclonal 3C10 (developed by C. Goodman and available from the Developmental Studies Hybridoma Bank) at 1:20. Nuclei were counterstained with DAPI. Embryos were stained using standard protocols ^39^.

### Cuticle Preparation

Embryos were collected and aged at 27°C until they were 16–22 hours old. Embryos were pipetted onto a slide and excess fluid was removed. Glacial acetic acid mixed 1:1 with Hoyer’s solution was added, covered with a coverslip, and allowed to dry for several days in an oven at 65°C for clearing. After 24 hours, the coverslips were weighted to flatten the preps. Cuticles were imaged on an upright Zeiss Axio Imager microscope with bright field illumination, and grayscale images were later inverted and oversaturated for increased contrast using Adobe Photoshop.

### Molecular characterization of protective indel mutations

To examine the molecular changes that conferred protection from dCas9-mediated overexpression and associated lethality, four genomic loci that include target sites for four functional sgRNAs (Figure S4) were amplified and sequenced. Single-fly genomic DNA preps were prepared using the solid tissue protocol of the *Quick*-DNA™ Miniprep Plus Kit (Zymo Research). In total, 2–3 μl of genomic DNA was used as a template in a 50-μL PCR reaction with Q5^®^ High-Fidelity 2X Master Mix (NEB, Ipswich, MA, USA). The following primers (Table S1) were used to amplify the loci with the corresponding sgRNA targets: 1001.S1 and 1001.S4 for hh; 1045.S1 and 1045.S4 for *wg*; 1045.S5 and 1045.S8 for *eve*; and 1045.S9 and 1045.S12 for *hid*. PCR products were loaded and run on an agarose gel, excised, and purified using a Zymoclean™ Gel DNA Recovery Kit (Zymo Research, Irvine, CA, USA) and were sequenced in both directions using Sanger sequencing at Retrogen Inc (San Diego, CA, USA). To characterize molecular changes at the targeted sites, AB1 sequence files were aligned against the corresponding reference sequences (downloaded from FlyBase release FB2019_3)^40^ in SnapGene® 4 and/or Benchling™.

### Gene Drive safety measures

All crosses using gene-drives genetics were performed in accordance to an Institutional Biosafety Committee-approved protocol from UCSD, in which full gene-drive experiments are performed in a high-security ACL2 barrier facility and split-drive experiments are performed in an ACL1 insectary in plastic vials that are autoclaved prior to being discarded in accordance with currently suggested guidelines for the laboratory confinement of gene-drive systems ^41,42^.

### Ethical conduct of research

We have complied with all relevant ethical regulations for animal testing and research and conformed to the UCSD institutionally approved biological use authorization protocol (BUA #R2401).

## Acknowledgements

We thank V. Kumar for help with library preparations. Sequencing was performed at the Millard and Muriel Jacobs Genetics and Genomics Laboratory at the California Institute of Technology. We thank N. Windbichler for sharing published sgRNA lines. We also thank Dr. N. Perrimon and B. Ewen-Campen for sharing published sgRNA expressing flies and plasmids. We thank J. Reinitz for sharing antibodies.

## Author Contributions

O.S.A. conceptualized the study. A.B, I.S., T.Y., J.L., I.A., J.M.M and M.E.P performed various genetic, molecular experiments, immunohistochemistry, bioinformatic, and mathematical modelling described in the study. All authors contributed to writing, analyzed the data, and approved the final manuscript.

## Data and Materials Availability

RNA sequencing data is available at NCBI SRA under accession number PRJNA578541. Fly strains engineered in this study to generate SPECIES will be available at Bloomington fly stock center with stock numbers listed in Fig S1. SPECIES lines will be made available upon request.

## List of Supplementary Materials

### Tables S1-S18

**Table S1.** Lethality and Rescue by targeting single genes

**Table S2.** Lethality and Rescue by targeting multiple genes

**Table S3.** Laying/survival experiments for SPECIES

**Table S4.** Interspecies crosses

**Table S5.** Embryo collections for RNA Sequencing

**Table S6.** RNAseq expression values for all genes and all 23 samples sequenced.

**Table S7.** TPM expression analysis for genes of interest.

**Table S8.** WTxA vs. A deseq2 Data

**Table S9.** WTxB vs. B deseq2 Data

**Table S10.** WTxC vs. C deseq2 Data

**Table S11.** WTxD vs. D deseq2 Data

**Table S12.** DEseq analysis comparing SPECIES outcrosses to SPECIES self crosses.

**Table S13.** Gene Drive Experimental Data

**Table S14.** Gene Drive Experimental Raw Data

**Table S15.** Primers used in this study.

**Table S16.** RNA Sequencing Statistical Data

**Table S17.** Population studies with associated fitness costs of SPECIES strains relative to WT

**Table S18.** Correlation between all RNA-seq samples.

**Table S19.** Outcrosses to Global Diversity Lines.

## SUPPLEMENTARY FIGURES

**Figure S1.**
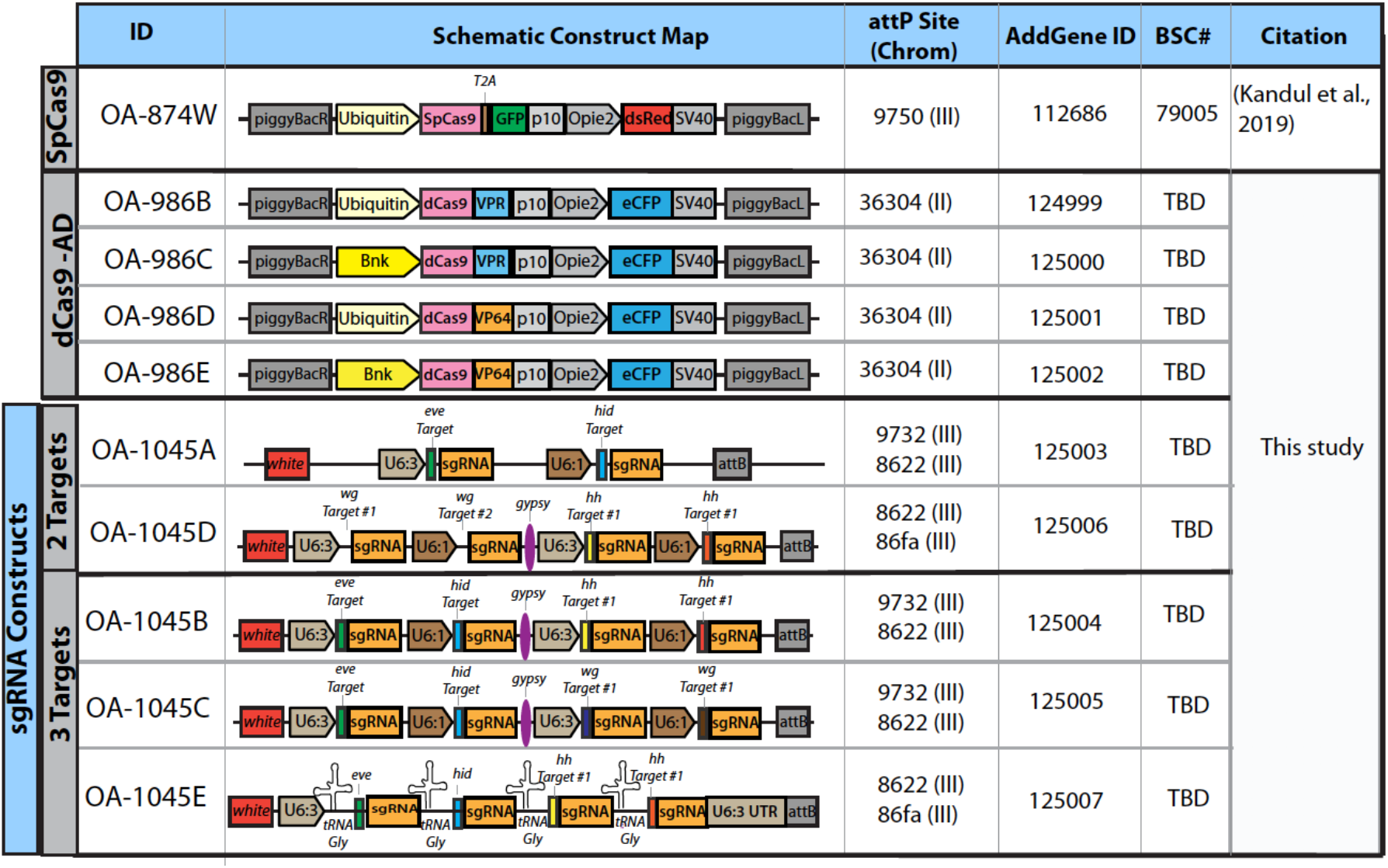
Constructs used in this study. A list of constructs used in this study, providing the construct ID, construct schematics, chromosomal insertion sites, Addgene ID number, Bloomington Stock number and citation.

**Figure S2.**
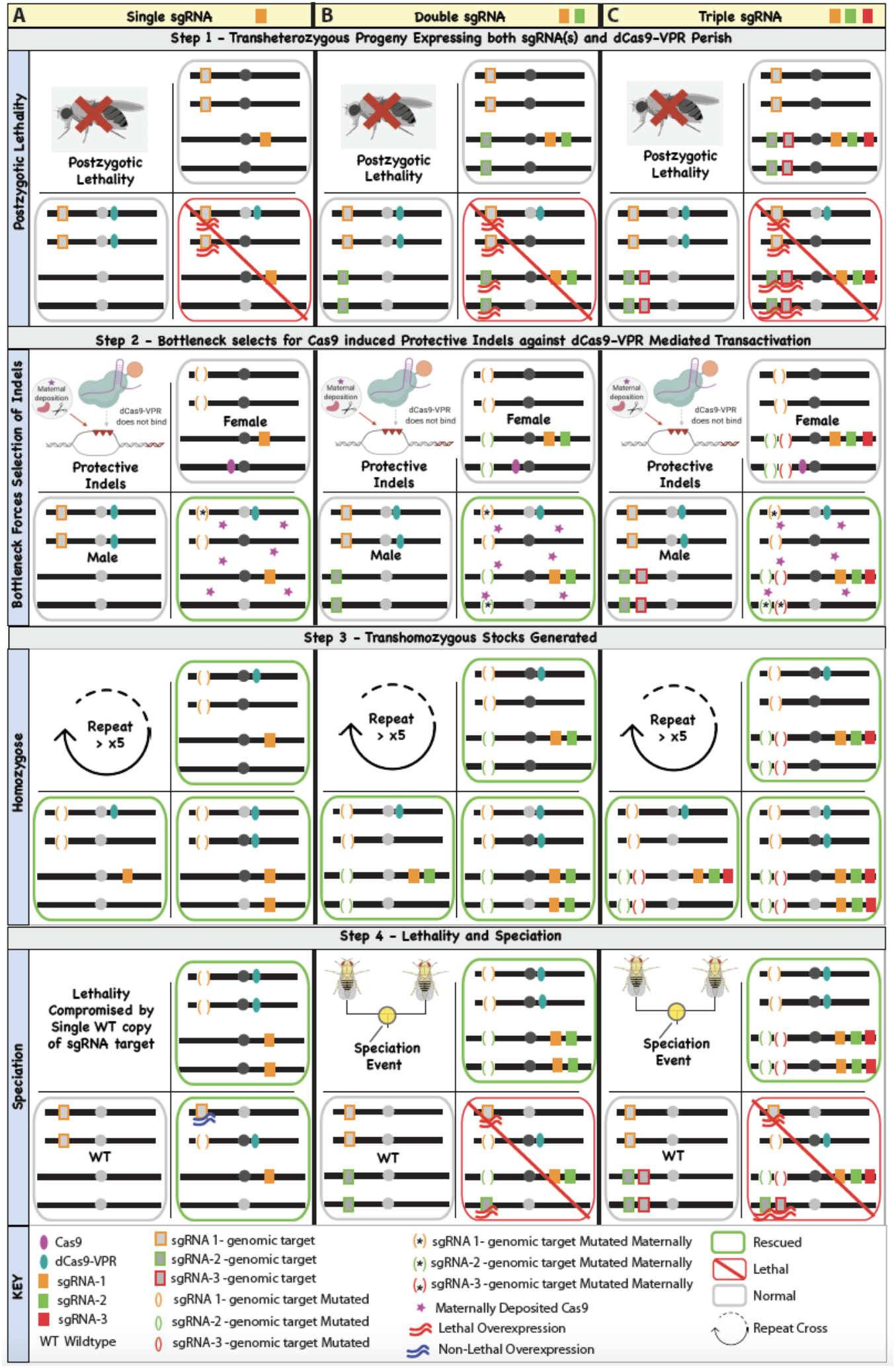
Schematic of the genetic crossing scheme used to engineer SPECIES. **(A)** Complete lethality (100%) was observed in transheterozygotes (dCas9/+; sgRNA/+) when an sgRNA was crossed to dCas9-VPR due to lethal overexpression (Step 1). To generate protective indels, the sgRNA was first crossed to Cas9, then transheterozygous (Cas9/+; sgRNA/+) females were crossed to dCas9-VPR males generating a bottleneck by which a small proportion of transheterozygotes (dCas9/+; sgRNA/+) survived due to protective indels generated by Cas9/sgRNA (Step 2). Surviving individuals (inheriting Cas9 protein maternally but lacking Cas9 as a gene) were inbred for many generations (>5) to generate homozygous stocks (Step 3). To assess lethality and speciation, homozygous stocks were bidirectionally outcrossed to WT. For a single sgRNA system, complete synthetic lethality and speciation was not observed due to the fact that one WT copy of the target promoter was not sufficient to induce lethal overexpression (A, Step 4). To overcome this issue, we multiplexed using either two sgRNAs **(B)**, or three sgRNAs **(C)**, and repeated steps 1-4 to engineer reproductively isolated synthetic species.

**Figure S3.**
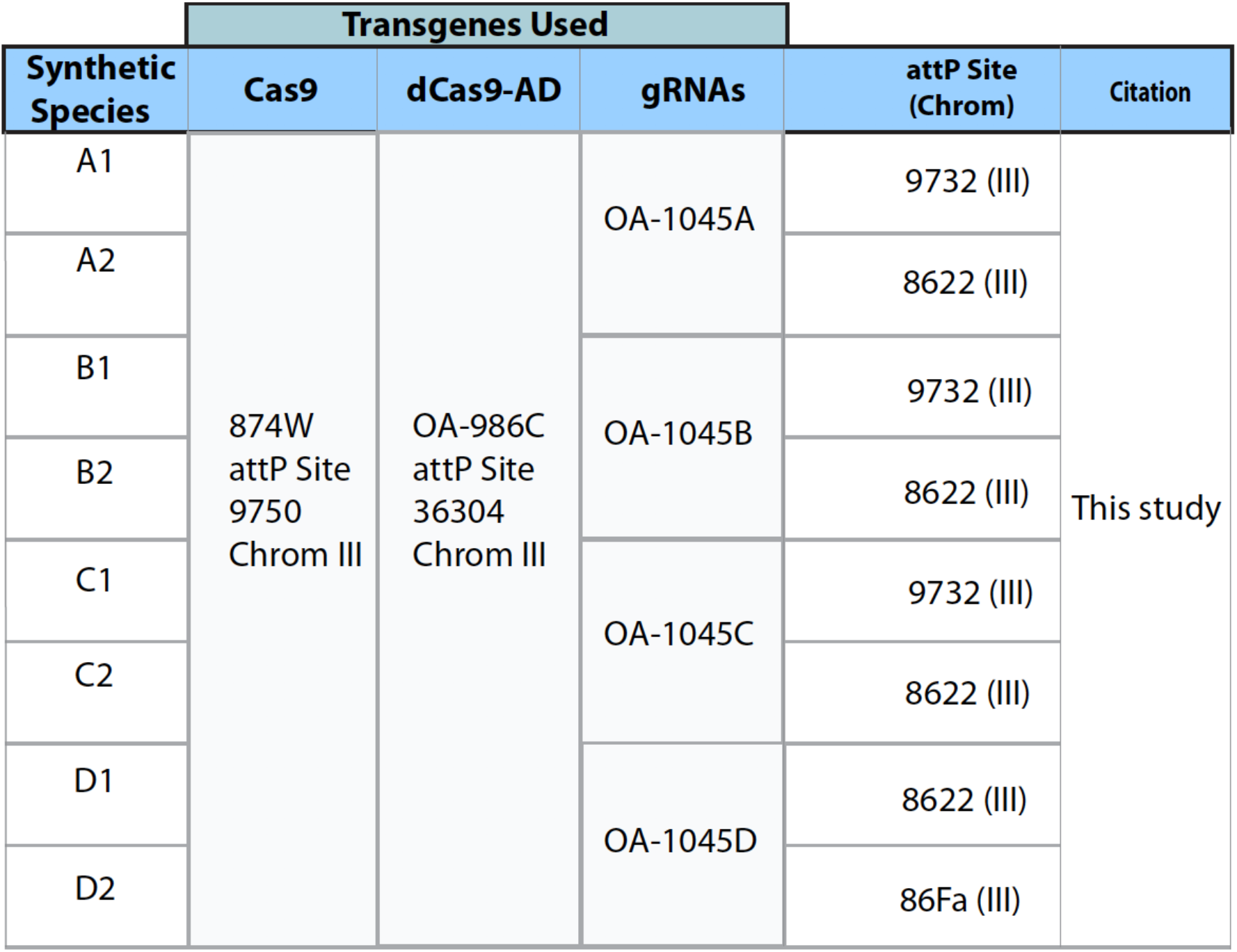
Generation of eight SPECIES. For each synthetic species (A1,A2,B1,B2,C1,C2,D1,D2) the transgene ID, and chromosomal insertion site are listed.

**Figure S4.**
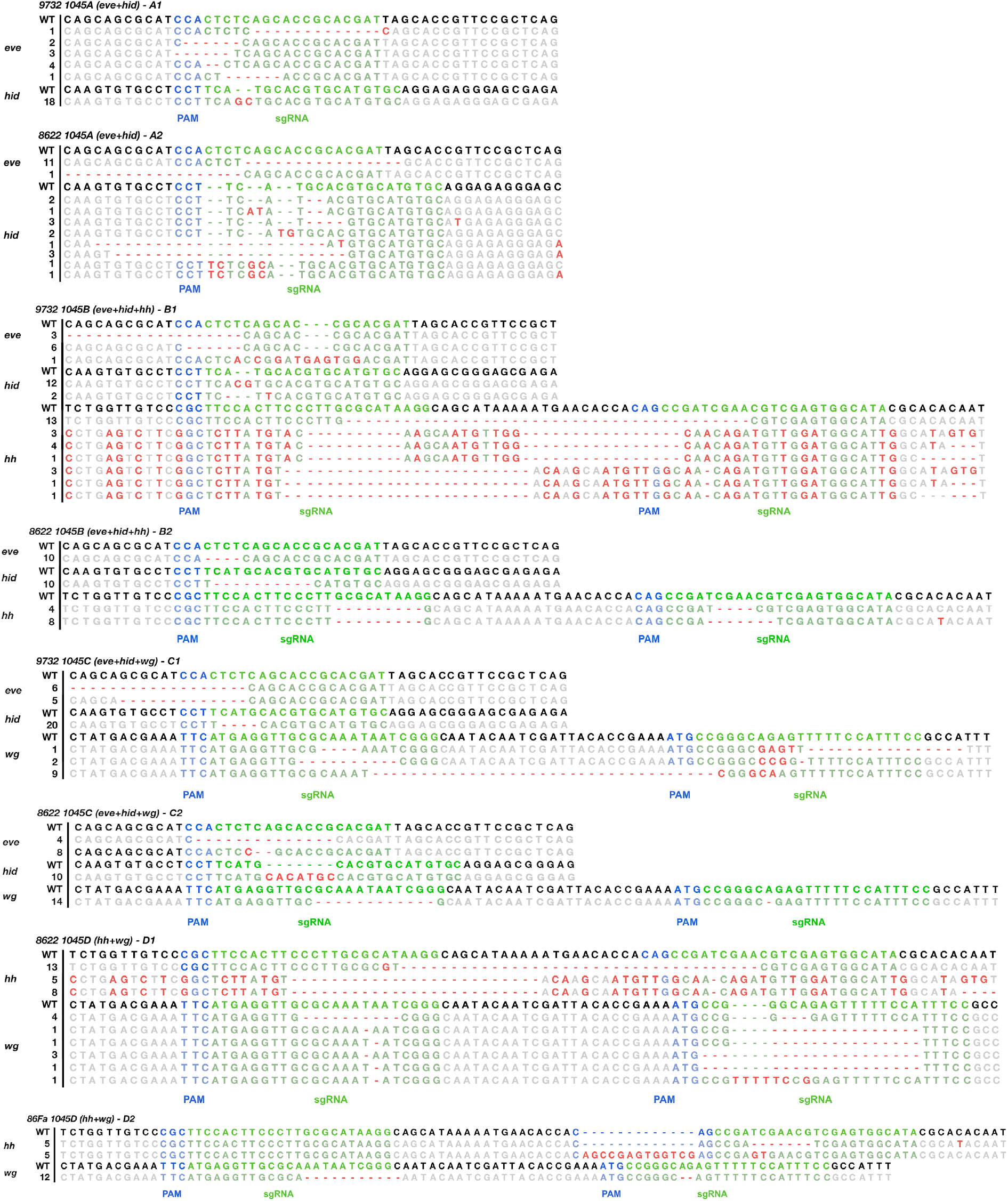
Molecular characterization of protective indel mutations. For the generation of each independent synthetic species (A1,A2,B1,B2,C1,C2,D1,D2) the gRNA target site was sanger sequenced and the indels were confirmed. Number to the left of each sequence indicates the number of individuals sequenced with this mutation.

**Figure S5.**
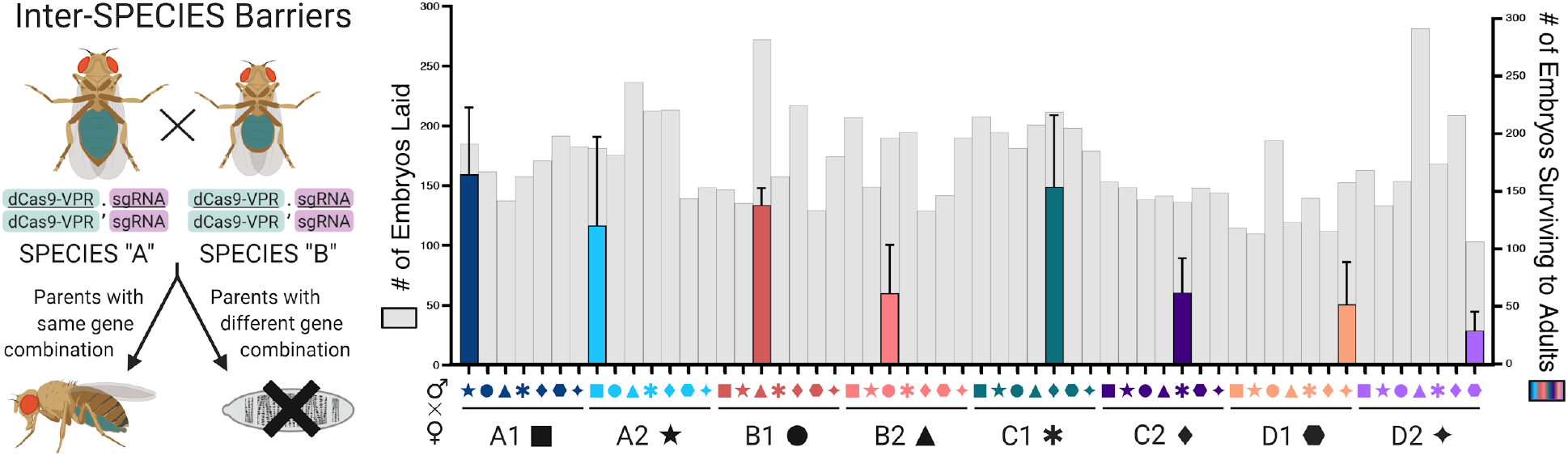
Reproductive isolation between double-homozygous speciated lines. Individuals from each SPECIES were crossed to one another to determine the extent of reproductive isolation between SPECIES (Table S4). The total # of embryos laid is plotted in grey on the left y-axis, while the total # of embryos surviving to adults is plotted on the right y-axis. All eight SPECIES were bidirectionally crossed to the remaining seven species in triplicate. The mean is plotted for each cross. Error bars indicate SD. In the graph, SPECIES “A” is listed by color and there is a symbol indicating SPECIES “B” in the cross (i.e., the star above the A1 group indicates the cross between an A1 female and an A2 male).

**Figure S6.**
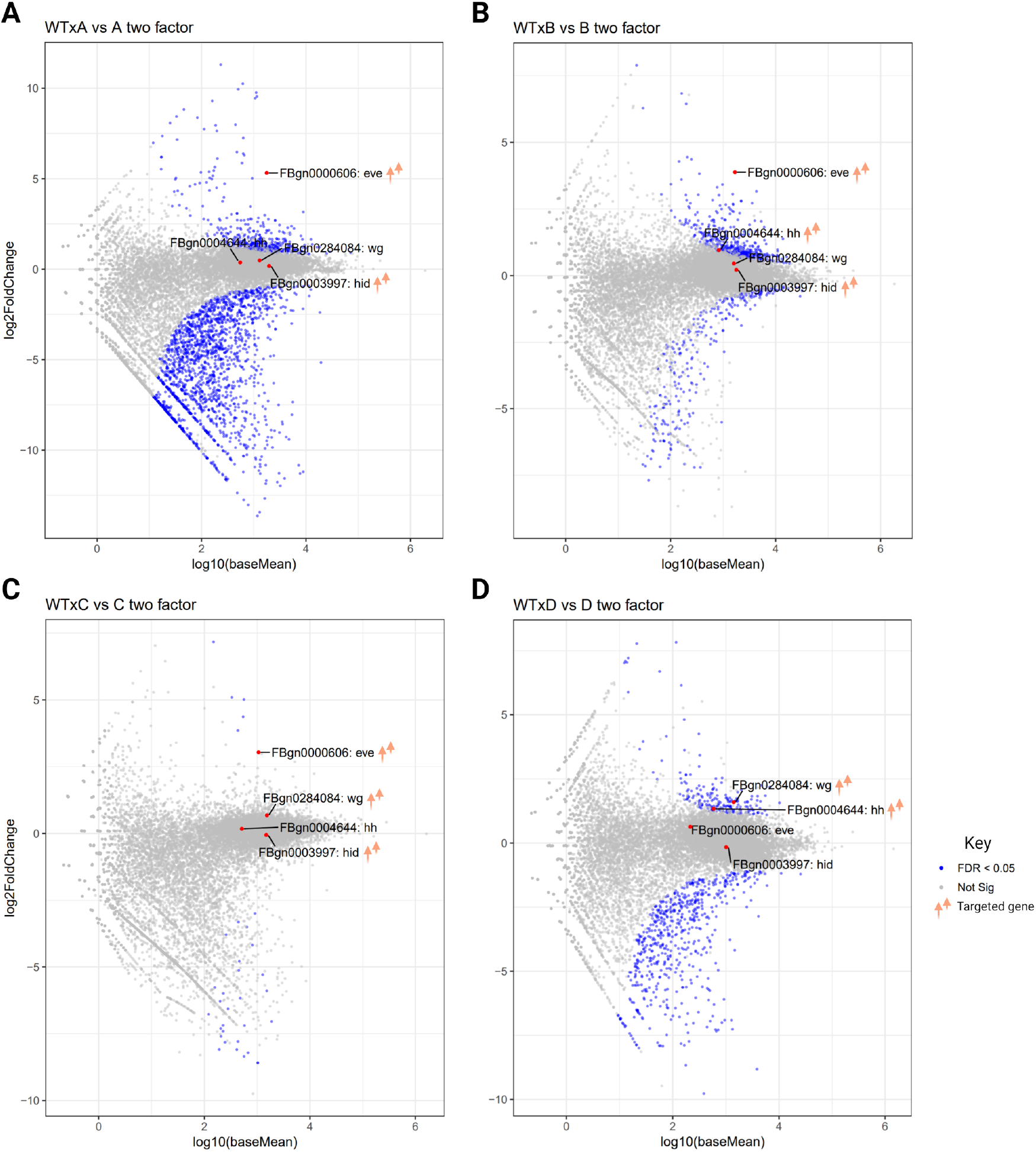
Two-Factor RNAseq comparisons. **(A)** Deseq comparisons between RNAseq samples WTXA1 (sample 8), WTXA2 (sample 9), A1XA1 (sample 10), A2XA2 (sample 11) observing target/non-target gene misexpression. **(B)**Deseq comparisons between RNAseq samples WTXB1 (sample 10), WTXB2 (sample 20), B1XB1 (sample 16) and B2XB2 (sample 22). **(C)** Deseq comparisons between RNAseq samples WTXC1 (sample 11), WTXC2 (sample 21), C1XC1(sample 17), and C2XC2 (sample 23). (D) Deseq comparisons between RNAseq samples WTD1 (sample 12), WTXD2(sample 13), D1XD1(sample 18), and D2XD2(sample 19). A full list of RNAseq sample IDs are listed in Table S5 and RNAseq data can be found in Tables S5-S12. Genes that are significantly misexpressed (FDR < 0.05) are colored blue.

**Figure S7.**
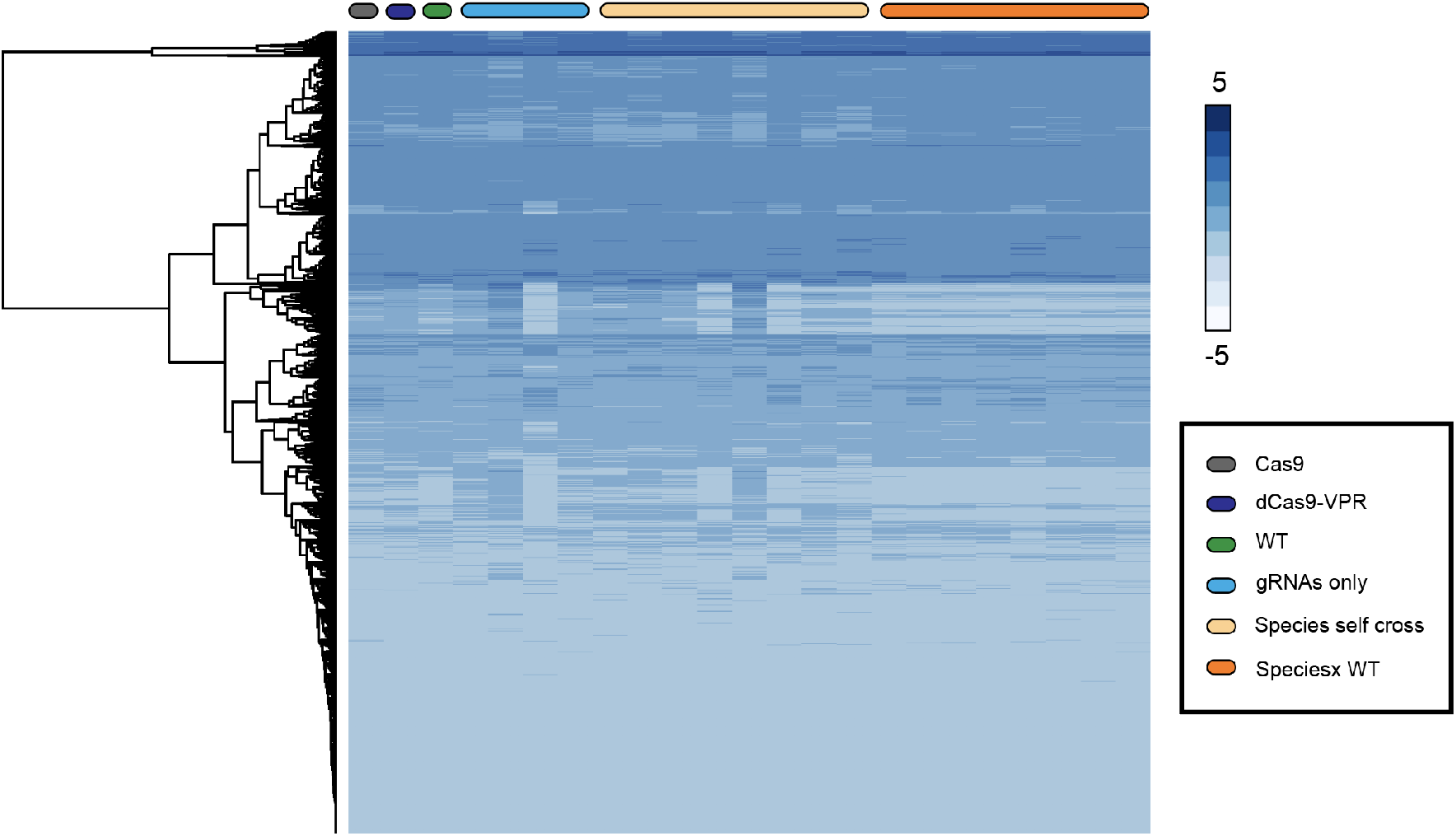
Heatmap of the RNAseq Data. Hierarchical clustering heat map of the RNAseq data (Table S6).

## SUPPLEMENTARY VIDEOS

**Video S1. Animation on engineering SPECIES.** A video explaining the process of how to generate SPECIES, covering CRISPR gene editing, the CRISPR-based transactivation, the self-cross and outcross experiment, potential applications, and significant advance. Embryos from twenty four independent genetic crosses consisting of each SPECIES (A1, A2, B1, B2, C1, C2, D1, D2) either self-crossed (SPECIES ♀ X SPECIES ♂), or bidirectionally out-crossed to WT (SPECIES♀ X WT♂ or SPECIES ♂ X WT♀) were laid on grape plates for 12 hours (# of embryos ranging from 27-135). Zoomed in images covering ~20% of the grape plate were taken once per day for 5 days. Embryo survival can be seen for each SPECIES when self-crossed but not when out-crossed to WT due to reproductive isolation.

**Table S15.**
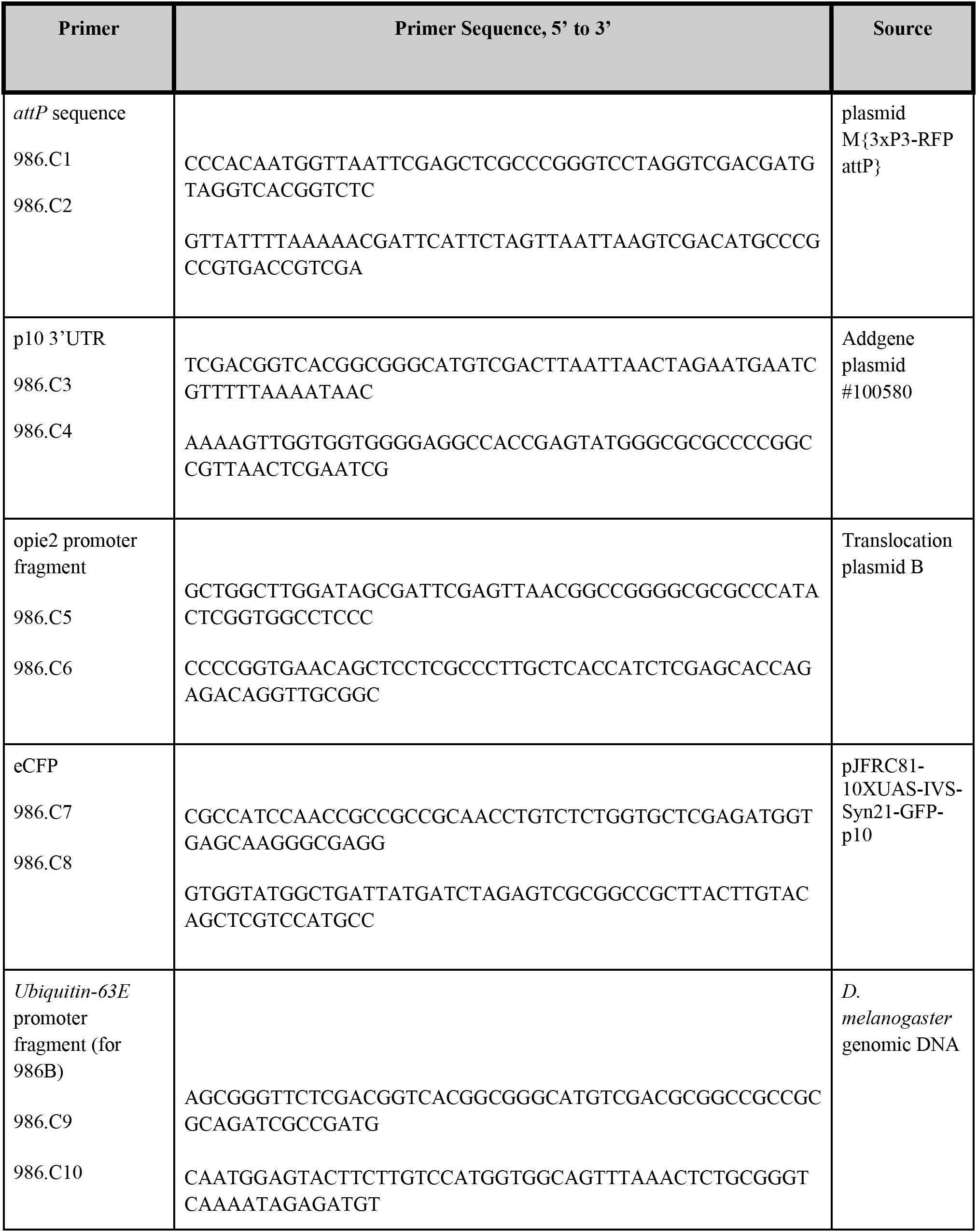
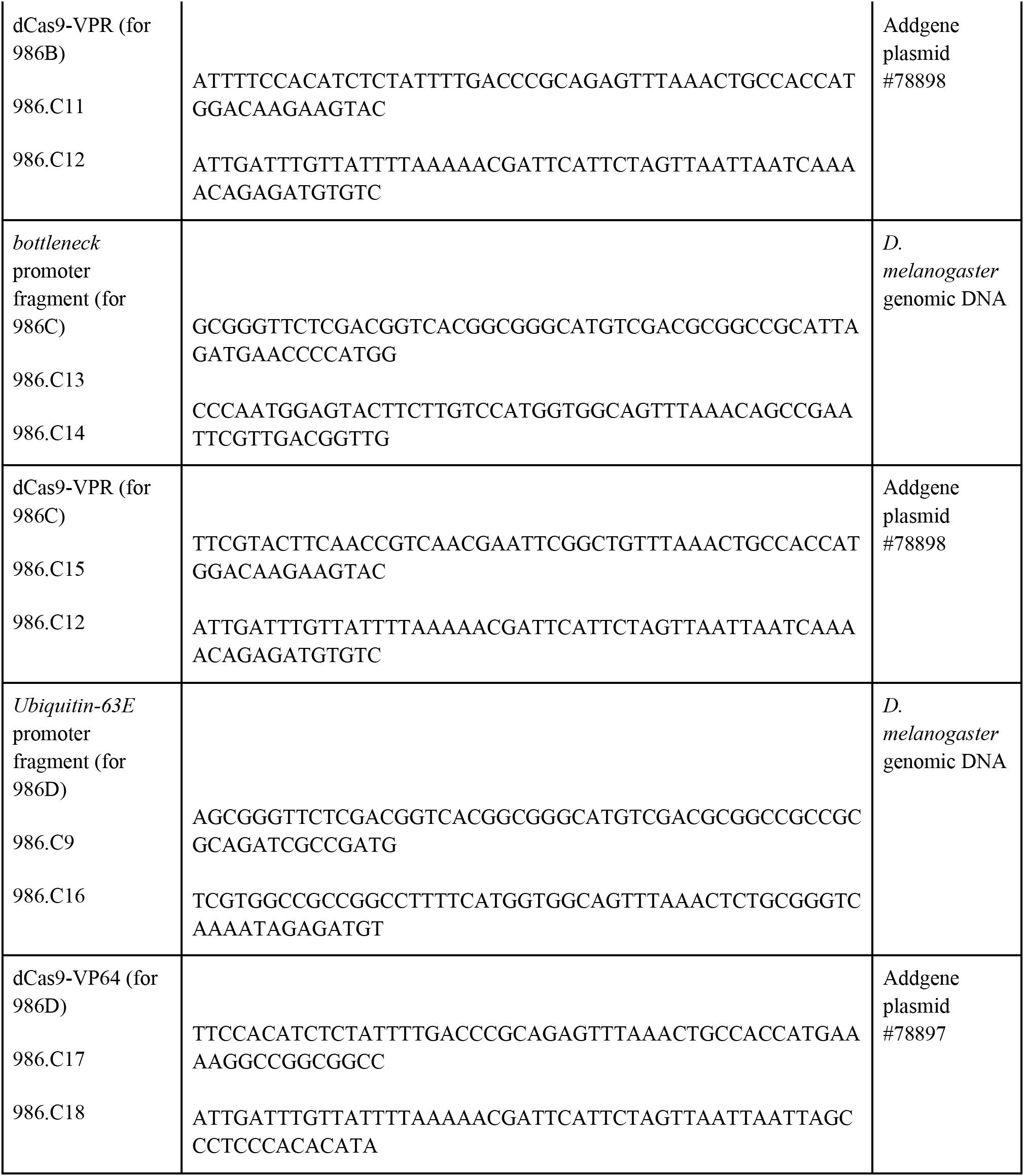
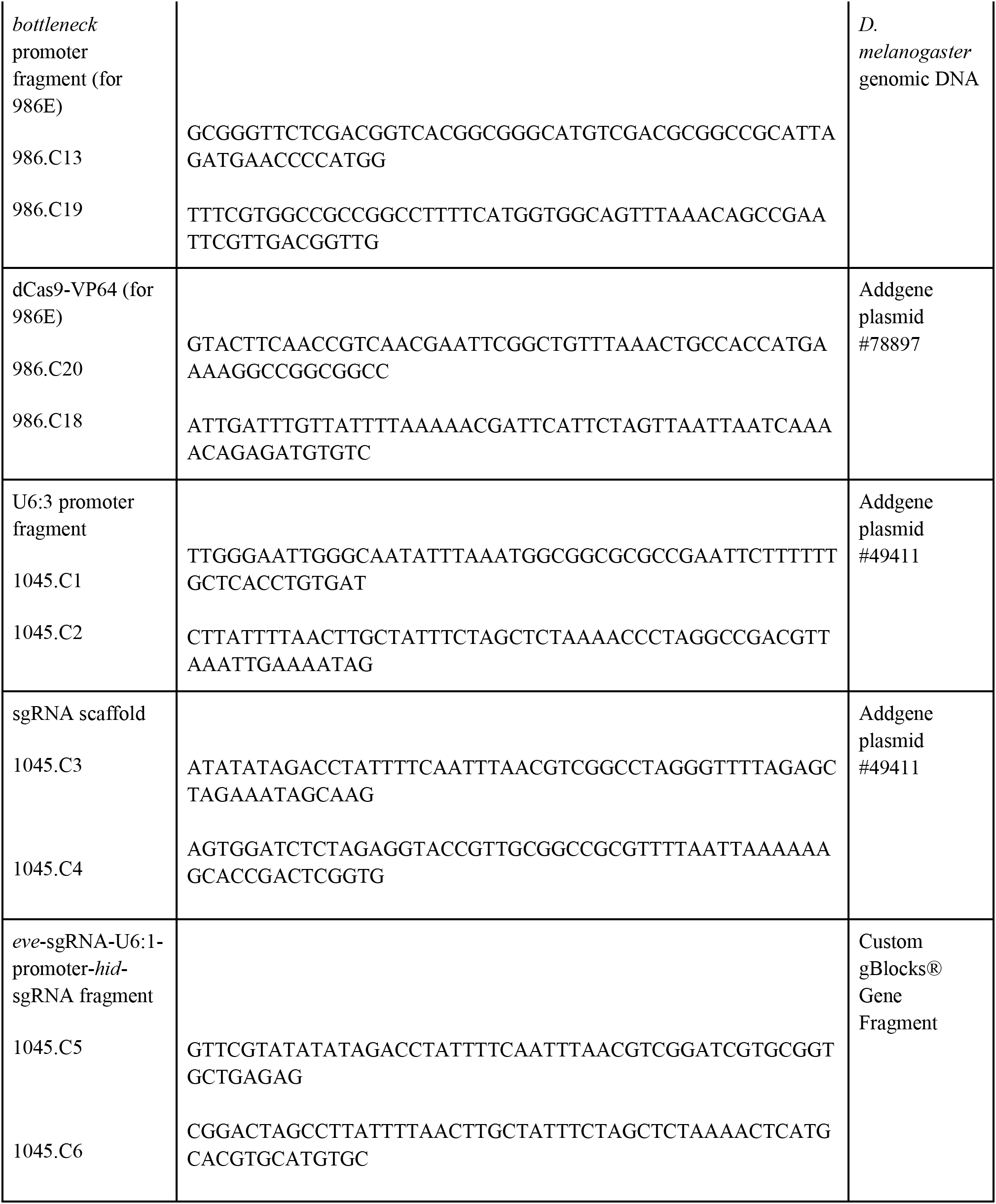
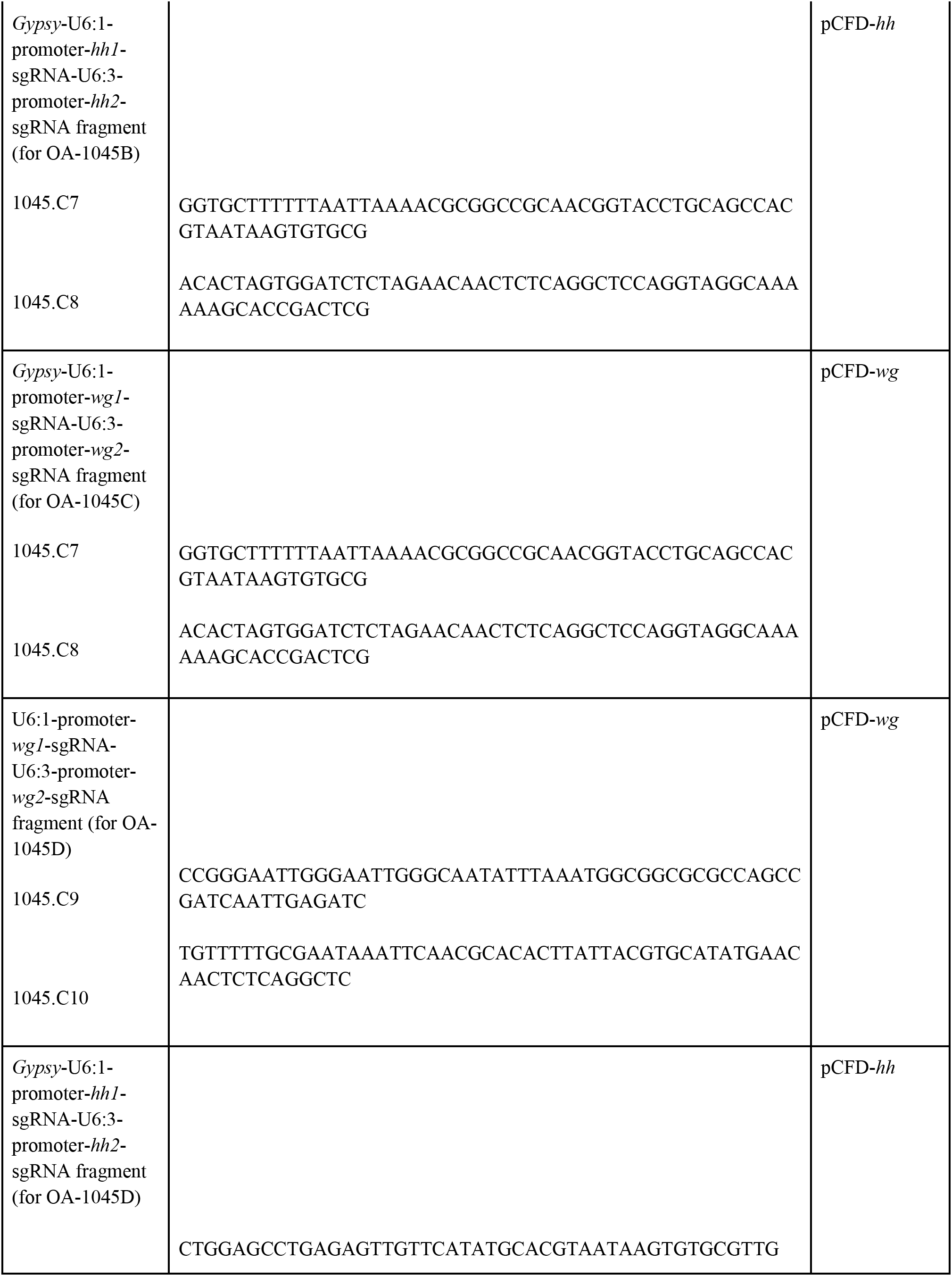
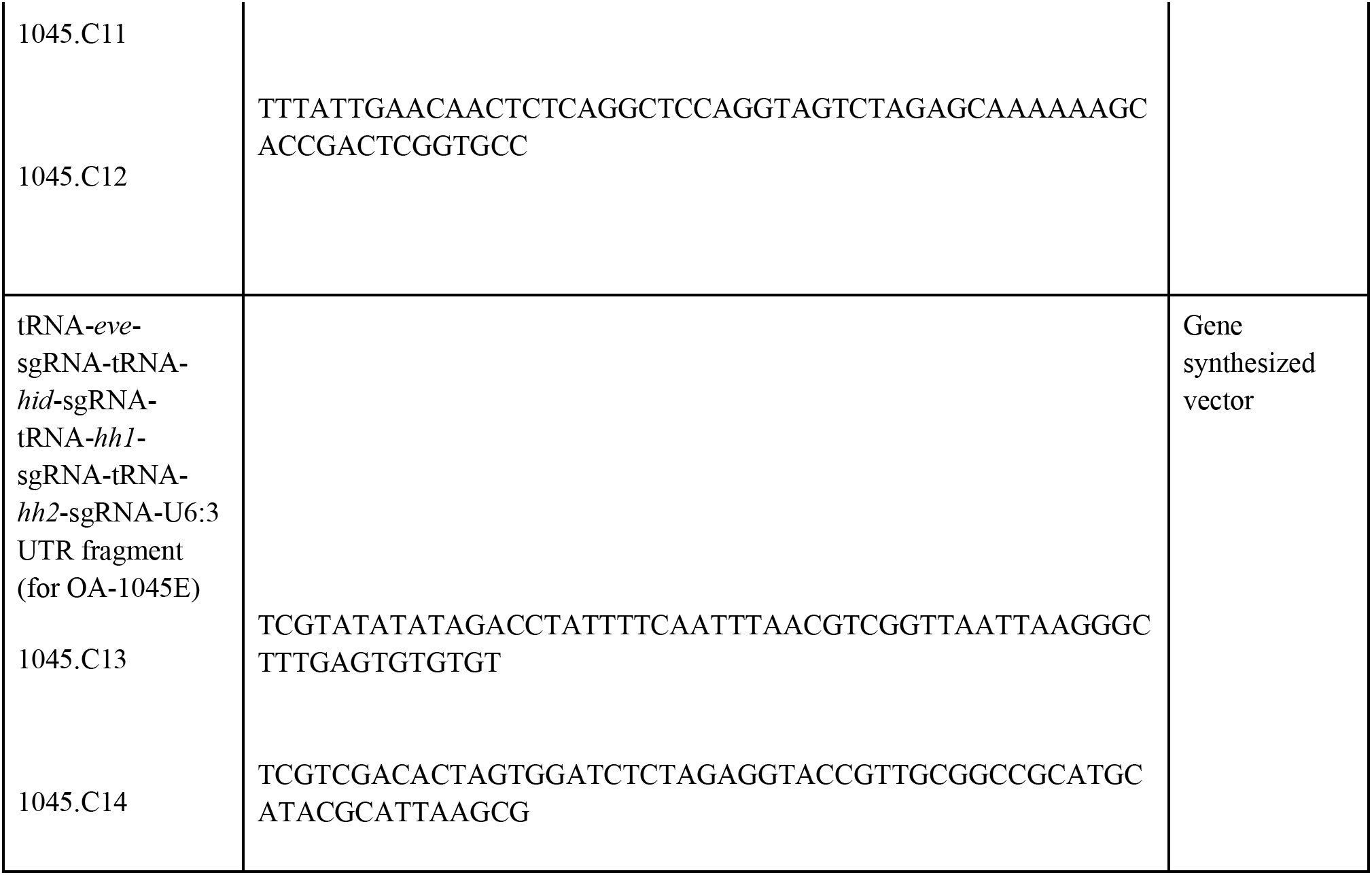
Primers used in this study.

